# Aperiodic EEG activity is a domain-specific marker of attentional aging in neurotypical adults

**DOI:** 10.64898/2026.07.22.739645

**Authors:** Christian Cazares, Alison Rigby, Leena Kang, Stephan Getzmann, Stefan Arnau, Edmund Wascher, Bradley Voytek

**Affiliations:** Department of Cognitive Science, University of California, San Diego, La Jolla, CA; Neurosciences Graduate Program, University of California, San Diego School of Medicine, La Jolla, CA; Leibniz Research Centre for Working Environment and Human Factors (IfADo), Dortmund, Germany; German Center for Mental Health (DZPG), Partner Site Bochum-Marburg, Bochum, Germany; Halıcıoğlu Data Science Institute, University of California, San Diego, La Jolla, CA, USA; Kavli Institute for Brain and Mind, University of California, San Diego, La Jolla, CA, USA

## Abstract

Aperiodic neural activity has been reliably shown to be reduced among older adults, and these reductions are associated with age-related cognitive decline. These results suggest that aperiodic activity, as measured by the spectral exponent, might be a viable electrophysiological biomarker for cognitive aging. However, this has not been tested longitudinally, within participants, during normal aging. Here, we used a longitudinal cohort to test whether changes in aperiodic activity between two sessions, approximately 5 years apart, track behavioral change across two domains: sustained attention (Psychomotor Vigilance Test) and executive function (modified Simon task). We found that aperiodic activity declined within individuals after the interval, independent of age at baseline. This within-person change tracked concurrent changes in reaction times during visual attention but was not associated with changes in any Simon task measure. Within a resampling framework, the aperiodic-PVT association was stable across cohort resamples and independently fitted partitions, though it was not confirmed in a small, frozen held-out set. These findings suggest the longitudinal decline in aperiodic activity specifically tracks sustained-attention processing speed rather than general cognitive aging, supporting its potential as a domain-specific biomarker of attentional aging in neurotypical populations.

## Introduction

In the United States, the adult population aged 65 and older is growing faster than any other age group^1^. This expanding share of the U.S. population is at the highest risk for age-related cognitive decline, Alzheimer’s disease, and other dementias, that may erode quality of life and threaten independent living^2–5^. As this population grows, we face an urgent need to identify objective, scalable neural markers that can detect cognitive change early and within individuals throughout their lifespan. In neurotypical adults, cognitive-behavioral functions that are compromised by aging include sustained attention^6^, information-processing speed^7^, and some aspects of executive function^8^. What remains unclear is how to index the neural processes underlying cognitive-behavioral changes in these domains via non-invasive biomarkers.

A promising approach is to leverage electrophysiological measures from non-invasive scalp electroencephalography (EEG), with one emerging candidate being aperiodic (broadband, non-oscillatory) neural activity^9–13^. This characteristic feature can be indexed by the aperiodic exponent, a measure that reflects the slope of the broadband 1/f-like background of the neural power spectrum in log-log space. This feature is physiologically distinct from the narrowband oscillations that have historically dominated cognitive EEG research^9,14^. In the search for mechanistically interpretable biomarkers of cognitive-behavioral aging, a growing literature links the aperiodic exponent to excitation/inhibition balance in cortical networks^15–17^, though there are likely many factors that contribute to it^18^. In neurotypical and compromised aging, this manifests as a decrease in the aperiodic exponent, indexing a shift toward excess excitation as a function of age-related loss of neuronal inhibition^10,19–26^. Such decreases in the aperiodic exponent can be visualized as the “flattening” of the spectral slope. Recent work across neural scales supports this interpretation, while also clarifying that the mapping between the exponent, aging, and E/I balance is not unconditional and varies by methodology and cell-type specificity^27–34^.

Cross-sectional studies have consistently shown a lower aperiodic exponent (i.e., a flatter spectral slope) in older compared with younger adults ^10,23,33–35^, and the aperiodic exponent has been linked to individual differences in cognitive performance across a range of domains, including attention and memory^16,34–38^. However, within-person age-related changes in aperiodic neural activity and their relationship to cognition and behavior throughout the lifespan remains largely underexplored. Thus far, longitudinal studies measuring the aperiodic exponent have mostly focused on developmental periods from early childhood through mid-adolescence, or in older adults during stroke recovery ^39–41^. One additional study confirmed that the aperiodic exponent decreases over a multi-year interval in healthy adults aged 20 to 70 years at baseline^24^, aligned with prior findings that were limited to shorter intervals of up to one year^44,45^.

However, it remains unknown whether within-person changes in the aperiodic exponent track concurrent changes in cognitive-behavioral decline in neurotypical adults across several years. And if such longitudinal coupling exists, it remains unclear whether it is general, consistent with a broad neural noise account of cognitive aging, or domain-specific, indexing specific cognitive processes. Cross-sectional evidence hints at domain specificity, with aperiodic measures being associated with age-related decline in episodic and visual short-term memory, but not working memory ^37,40^. Closing this gap requires within-subject longitudinal analyses that pair changes in the aperiodic exponent with cognitive-behavioral measures across different domains, both to validate the exponent as a biomarker of within-person cognitive change and to specify the cognitive processes it tracks.

Here we leveraged a multi-year, longitudinal dataset from a cohort of neurotypical adults, sourcing data from the Dortmund Vital Study^46,47^, that were administered task-free scalp EEG before and after cognitive-behavioral assessments at two timepoints approximately 5 years apart. To dissociate domain-general from domain-specific effects over this period, we analyzed two cognitive measures: the Psychomotor Vigilance Test (PVT), an index of sustained attention and processing speed ^48,49^, and a modified version of the Simon task that served as an index of executive function and interference control^50,51^. We first established that the aperiodic exponent declined within individuals over the approximately five year time interval (consistent with a parallel report in a subset of this cohort^24^), independent of age at baseline, alongside slowing of PVT reaction time over the same interval. Critically, within-person change in the aperiodic exponent tracked concurrent change in PVT reaction time after controlling for baseline age, baseline exponent, baseline performance, and sex. This effect was observed only when using aperiodic exponent measures from EEG collected before, but not after, the cognitive-behavioral assessments. In contrast, exponent change did not track longitudinal change in any Simon measure. Together, these results suggest that the longitudinal decline in the aperiodic exponent, measured from task-free scalp EEG, may serve as a domain-specific marker of attentional aging rather than a general index of cognitive aging in neurotypical populations.

## Results

### The aperiodic exponent declines in neurotypical adults

We analyzed a dataset from 276 neurotypical adults with task-free EEG measured before and after a 2-hour block of cognitive-behavioral testing across two timepoints that were an average of 5 years apart (5.3 ± 0.8 years, mean ± SD; 166 females). At both timepoints, three minutes of EEG was collected under two conditions, eyes closed and eyes open, before (pre-task) and after (post-task) completing the PVT and Simon tasks (**Fig. 1A**). Data from the full cohort was split *a priori* into a training set (n = 222; 131 females) and a held-out test set (n = 54; 35 females) reserved for the later predictive analyses, stratified to match in baseline age (Mann-Whitney U = 5,864, p = 0.80) and sex (χ^2^(1) = 0.39, p = 0.53) (**Fig. 1B**) (**Table 1**). All remaining analyses used the training set until predictive analyses were performed.

**Figure 1.**
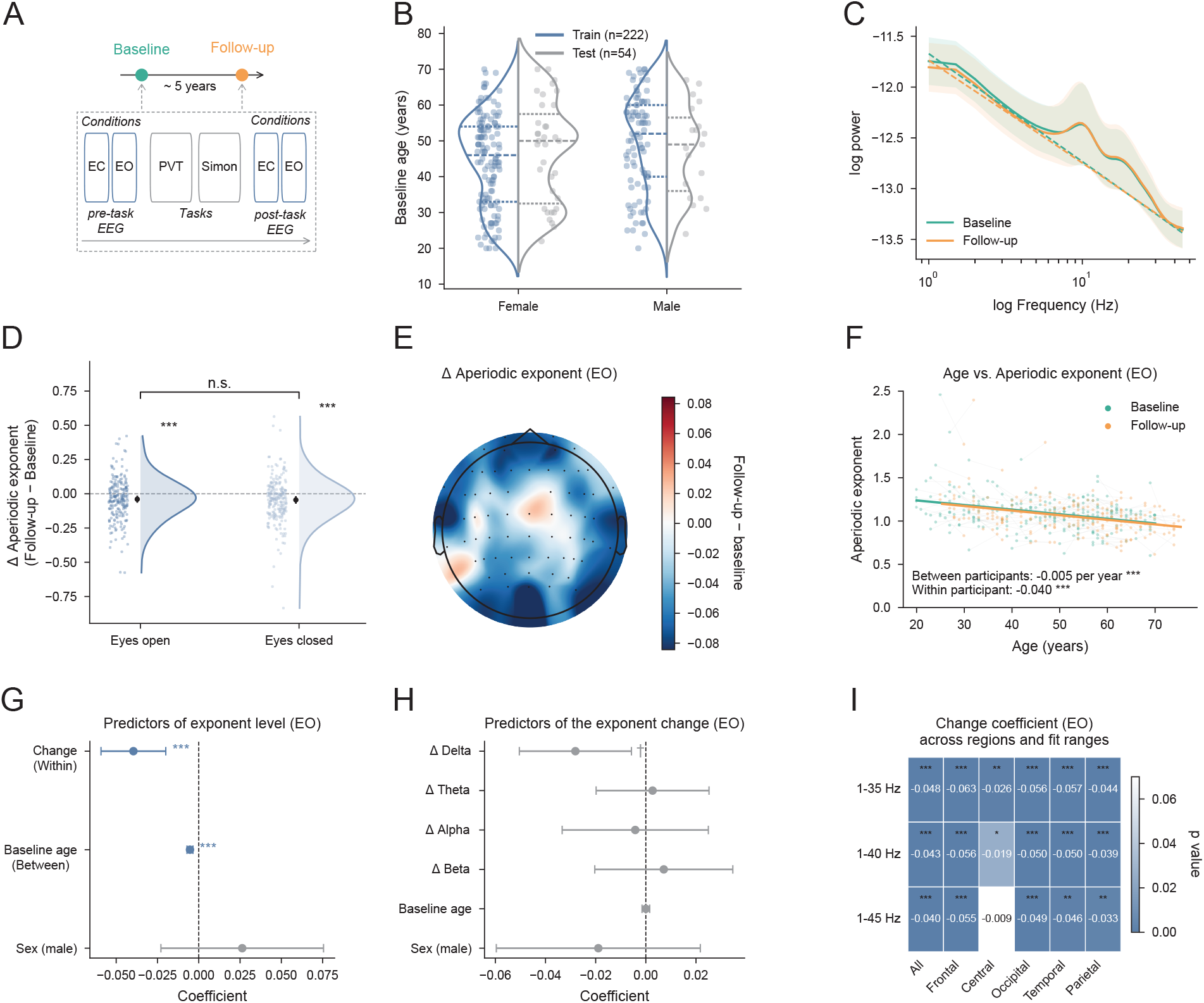
The aperiodic exponent declines within neurotypical adults. (A) Study design. Sessions were identical at baseline and at the approximately five-year follow-up: pre-task EEG (eyes closed then eyes open), the PVT and Simon tasks (in addition to three additional tasks not analyzed in this study), then post-task EEG. (B) Baseline age distribution by sex for the training (blue, n = 222) and held-out test (grey, n = 54) sets. Split violins mark quartiles with individual participants overlaid. The sets did not differ in baseline age (Mann-Whitney U = 5,864, p = 0.80) or sex (chi-squared test, χ2(1) = 0.39, p = 0.53). (C) Group power spectra plotted in log-log space at baseline (teal) and follow-up (orange) with the aperiodic fit overlaid (dashed). Lines and shading show the mean and ±1 standard deviation for all electrodes across participants. (D) Within-person exponent change (follow-up minus baseline) under eyes-open and eyes-closed conditions. Clouds show the participant distribution and the point and whisker show the mean and its 95% confidence interval. The exponent declined in both conditions (eyes open mean = -0.040; eyes closed mean = -0.045; both two-sided sign-flip permutation test, 10,000 permutations, p < 0.001) and the decline did not differ between conditions (sign-flip permutation, paired t(221) = 0.46, p = 0.64). (E) Scalp topography of the mean exponent change across 64 channels (eyes-open condition). (F) Exponent against age at baseline (teal) and follow-up (orange) with mixed-effects lines overlaid (all electrodes mean, eyes-open condition). Line slope is the between-person age gradient and the offset between lines indicates the within-person change. (G) Mixed-effects coefficients for exponent level: within-person change, baseline age, and sex (all electrodes mean, eyes-open condition). Error bars are 95% confidence intervals with significant terms in blue and non-significant in grey. (H) Coefficients relating concurrent band-power change to exponent change (all electrodes mean, eyes-open condition). No band predicted the decline (ordinary least-squares regression, Benjamini-Hochberg FDR; all FDR-corrected p > 0.05). (I) Within-person change coefficient (linear mixed-effects model) across three spectral parameterization fitting ranges and six electrode groups; shading denotes the p-value and asterisks mark significance (all p < 0.05 except for the central region at 1-45 Hz). n = 222 (training set) in C to I; n = 276 in B. † < 0.07, *p < 0.05, **p < 0.01, ***p < 0.001.

**Table 1.** Cohort characteristics for the training and held-out test splits. Demographic and follow-up characteristics for the training split (n = 222), the a priori held-out test split (n = 54) and the full cohort (n = 276). N is the number of participants; Female is given as count (percentage); baseline age and the follow-up interval are given as mean (standard deviation, SD) with the observed range, where the follow-up interval is the per-participant elapsed time in years between the baseline and follow-up sessions. the within-person aperiodic decline was a stable neural measure for the behavioral analyses that followed.

|  | Training Split | Held Out Test Split | Full cohort |
| --- | --- | --- | --- |
| *N* | 222 | 54 | 276 |
| Female, n (%) | 131 (59.0%) | 35 (64.8%) | 166 (60.1%) |
| Baseline age, mean (SD) | 46.581 (13.349) | 47.185 (13.631) | 46.699 (13.382) |
| Baseline age, range | 20–70 | 22–70 | 20–70 |
| Follow-up interval (yr), mean (SD) | 5.365 (0.771) | 5.111 (0.744) | 5.315 (0.771) |
| Follow-up interval (yr), range | 4–8 | 4–7 | 4–8 |

Across baseline and follow-up sessions, task-free EEG power spectra followed the canonical aperiodic (1/f-like) relationship (**Fig. 1C**). The aperiodic component fitted to the participant-averaged spectra for all electrodes was ‘flatter’ at follow-up, consistent with a reduced aperiodic exponent. The exponent declined in both task-free conditions (eyes open mean change = -0.040, 95% CI [-0.059, -0.020], permutation sign-flip test p < 0.001, d_z_ = -0.27; eyes closed mean change = -0.045, 95% CI [-0.066, -0.023], p < 0.001, d_z_ = -0.28) (**Fig. 1D**). This decline did not differ between eyes-open and eyes-closed conditions (sign-flip permutation test, paired t(221) = 0.46, p = 0.64, d_z_ = 0.03). Due to this lack of difference, we focused remaining analyses on the eyes-open condition unless stated otherwise. Visually, this aperiodic exponent decline was spatially distributed across the scalp (mean change = -0.04) (**Fig. 1E**).

To separate cross-sectional age differences from longitudinal change, we fitted a linear mixed-effects model of the exponent. Older ages at baseline were associated with a lower aperiodic exponent between individuals (β = -0.0053 per year, 95% CI [-0.0071, -0.0035], p < 0.001), and independently of this gradient, the aperiodic exponent declined within individuals (β = -0.040, 95% CI [-0.059, -0.020], p < 0.001), with no sex effect (β = +0.026, 95% CI [-0.023, 0.076], p = 0.29) (**Fig. 1F, G**) (see **Supplementary Table 1**). To test if the exponent decline was driven by concurrent shifts in oscillatory power, we regressed the within-person exponent change on the concurrent within-person changes in delta (1-4 Hz), theta (4-7 Hz), alpha (7-13 Hz), and beta (13-24 Hz) band aperiodic adjusted power, together with age at baseline and sex. No band-power change predicted the observed exponent changes (all FDR-corrected p > 0.05) (**Fig. 1H**), indicating that the decline was not merely a by-product of changes in oscillatory-power. To confirm that the mean exponent decline was robust to spectral parameterization fitting ranges, we refitted the within-person model across three spectral fitting ranges (1-35, 1-40, 1-45 Hz) and six regional electrode groups (**Fig. 1I**). The cross-session change coefficient remained largely negative (p < 0.05) with a single exception (central region at 1-45 Hz), suggesting that

### Attentional reaction time slows in neurotypical adults

We next characterized the aging-related change in sustained attention and processing speed as assayed by the PVT (**Fig. 2A**). In a linear mixed-effects model, mean reaction time rose across the ten-minute task and this time-on-task increase did not differ between sessions (*bin*, β = 7.9 ms per 2-min bin, 95% CI [6.48, 9.42], p < 0.001; *bin x session*, β = 0.80, 95% CI [-1.27, 2.88], p = 0.45) (**Fig. 2B**). Mean reaction time increased from baseline to follow-up (mean change = 10.5 ms, 95% CI [3.08, 17.98]; sign-flip permutation test, paired t(221) = 2.76, p = 0.006, d_z_ = 0.19) (**Fig. 2C**), whereas neither reaction time variability nor response accuracy changed over the same interval (sign-flip permutation test p = 0.45 and p = 0.38, respectively) (**Supplementary Fig. 1A, B**). Linear mixed-effects models of each measure (*measure ∼ session + age + sex + (1 | participant)*) confirmed these within-person changes and separated them from the between-person age gradient. Cross-sectionally, older baseline age was associated with faster baseline reaction time, opposite to the within-person slowing (β = -0.529 ms per year, 95% CI [-1.02, -0.04], p = 0.034) (**Fig. 2D, E**) (see **Supplementary Table 2**).

**Figure 2.**
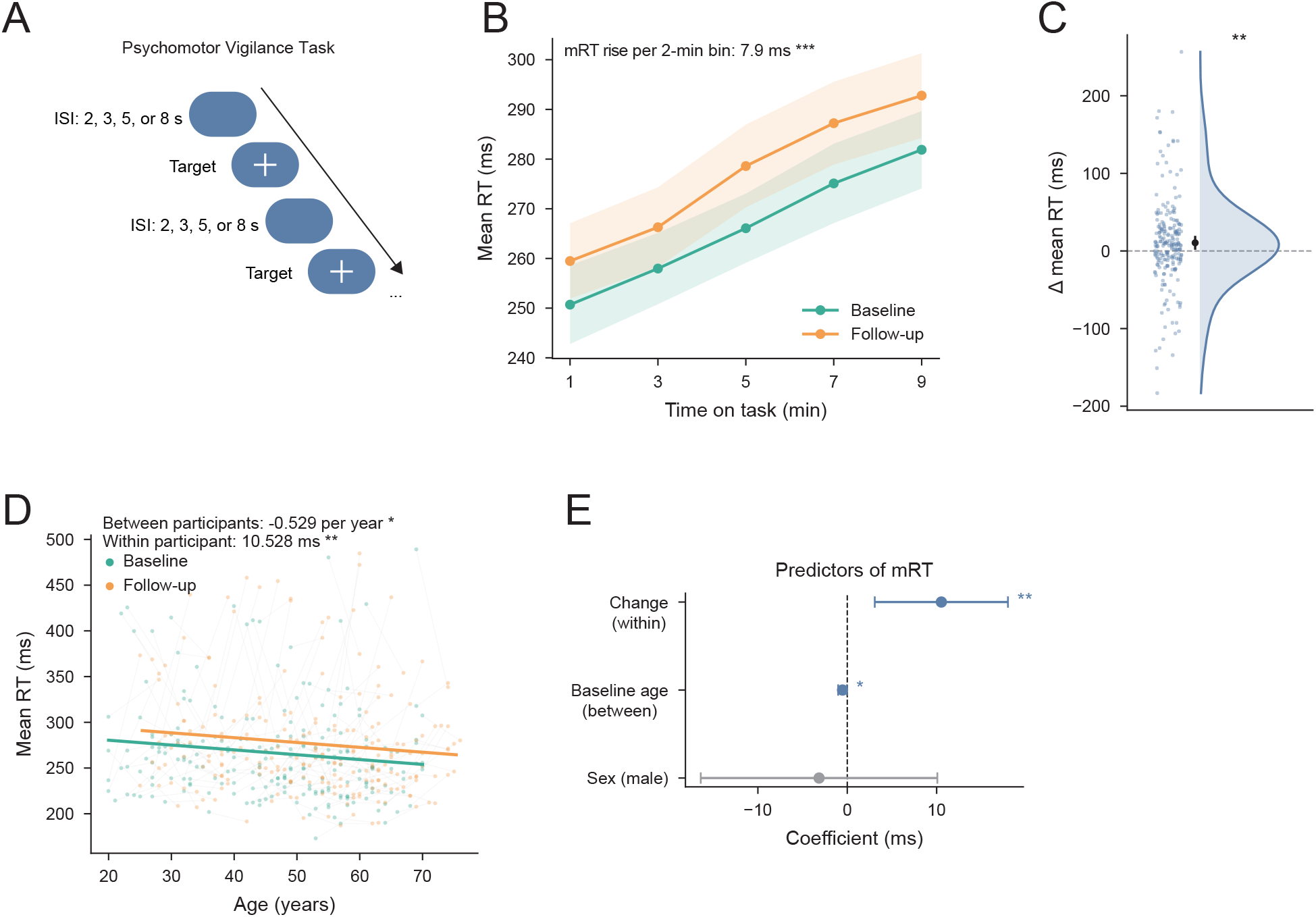
Reaction time in the PVT slows within neurotypical adults. (A) Schematic of the psychomotor vigilance task. After a variable interstimulus interval (ISI) of 2, 3, 5 or 8 s, a target (“+”) appeared and participants were instructed to respond to its onset with a button press as quickly as possible. (B) Mean reaction time across the ten-minute task in two-minute bins at baseline (teal) and follow-up (orange). Points and shading show the mean and 95% confidence interval. In a linear mixed-effects model, reaction time rose with time-on-task (β = 7.9 ms per bin, p < 0.001) and this did not differ between sessions (session x bin interaction p = 0.45). (C) Within-person change in mean reaction time (follow-up minus baseline). Clouds show the participant distribution and the point and whisker show the mean and its 95% confidence interval. Reaction time increased (mean = 10.5 ms, two-sided sign-flip permutation test, 10,000 permutations, p = 0.006). (D) Mean reaction time against age at both sessions with mixed-effects lines; older baseline age was associated with faster baseline reaction time (linear mixed-effects model, between-person β = -0.53 ms per year, p = 0.034), opposite to the within-person slowing. (E) Coefficients for mean reaction time from a linear mixed-effects model: within-person change, baseline age, sex. Error bars denote 95% confidence intervals. Significant terms in blue and non-significant in grey. n = 222 (training set) in B to E. *p < 0.05, **p < 0.01, ***p < 0.001.

Because change scores are subject to regression to the mean, we modelled the change in mean reaction time directly, accounting for baseline mean reaction time, baseline age, and follow-up duration influenced it. Baseline mRT predicted the change (β = -0.44 ms per ms baseline RT, 95% CI [-0.57, -0.32], p < 0.001, adj. R^2^ = 0.17), with slower participants at baseline tending to become faster (**Supplementary Fig. 1C**). The mean increase nonetheless survived this adjustment (intercept at mean baseline = 10.5 ms, 95% CI [3.70, 17.36], p < 0.001). After residualizing the change on baseline performance and follow-up interval, neither baseline age (β = -0.23 ms per year, 95% CI [-0.75, 0.30], p = 0.40) nor follow-up interval (β = 3.73 ms per year, 95% CI [-5.34, 12.79], p = 0.42) predicted its magnitude (**Supplementary Fig. 1D**), suggesting a cohort-wide, within-person slowing rather than an effect concentrated in older participants or moderated by any specific time-interval between sessions.

### The aperiodic exponent tracks concurrent attentional reaction time slowing

We next investigated whether the aperiodic exponent was correlated with the observed PVT mean reaction time increase. Across the pooled baseline and follow-up sessions, mean reaction time did not correlate with the exponent (β = 15 ms per unit, 95% CI [-24.52, 54.52], p = 0.46, baseline r = 0.1, follow-up r = -0.04), although the slope relating the exponent to reaction time grew more negative at follow-up (*exponent x session*, β = -41.5 ms, 95% CI [-82.95, -0.09], p = 0.0495) (**Fig. 3A**). Within individuals, participants with a greater decline in the exponent showed a greater increase in mean reaction time (r = -0.17, p = 0.010) (**Fig. 3B**), and this relationship was driven by the exponent change rather than the baseline exponent (r = 0.027, p = 0.69) (**Fig. 3C**). Both bivariate relationships remained when assessed using a multivariable model that included the change and the baseline exponent simultaneously, together with baseline age, baseline reaction time, sex and the follow-up interval. In this model, each one-unit decrease in the exponent corresponded to a 91 ms increase in reaction time (*Δexponent*, β = -91.11 ms, 95% CI [-142.63, -39.59], p < 0.001, FDR-corrected p = 0.008), whereas the baseline exponent remained non-significant (β = -11.9 ms, 95% CI [-49.37, 25.48], p = 0.53; adjusted R^2^ = 0.21) (**Fig. 3D**). This relationship was not moderated by baseline age (*Δexponent x baseline age*, β = -0.81, 95% CI [-4.02, 2.40], p = 0.62) and did not depend on the specific follow-up time interval (β = 4.1 ms per year, 95% CI [-4.80, 12.99], p = 0.37), suggesting a cohort-wide, within-person effect. Baseline reaction time was negatively related to its own change (β = -0.47, 95% CI [-0.59, -0.34], p < 0.001) consistent with regression to the mean, but the exponent coupling held over and above this dependence.

**Figure 3.**
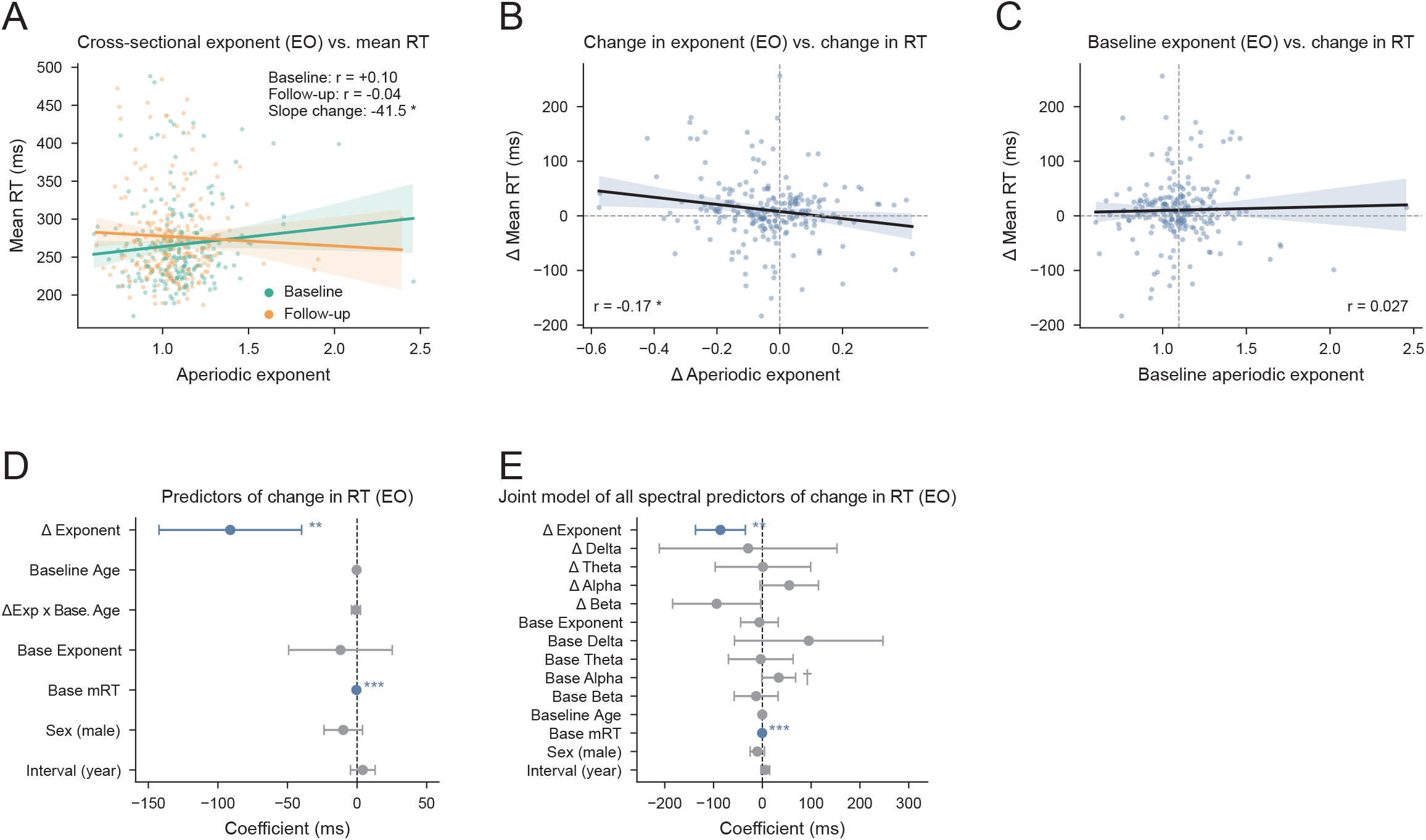
Within-person aperiodic exponent decline tracks concurrent reaction time slowing in the PVT. (A) Mean reaction time against the aperiodic exponent across pooled sessions at baseline (teal) and follow-up (orange). Lines and shading show the fit and 95% confidence interval from cluster-robust ordinary least-squares regression. The exponent did not relate to reaction time overall (β = 15, p = 0.46), though the slope grew more negative at follow-up (exponent x session β = -41.5 ms, p = 0.0495). (B) Within-person reaction time change against within-person exponent change. Participants with greater exponent decline slowed more (Pearson r = -0.17, p = 0.010). (C) Reaction time change against the baseline exponent did not show a relationship (Pearson r = 0.027, p = 0.69). (D) Coefficients from the full coupling model of reaction time change (ordinary least-squares with covariates: exponent change, baseline age, their interaction, baseline exponent, baseline mean reaction time, sex, follow-up interval). Each unit of exponent decline corresponded to a 91 ms increase in reaction time (exponent change β = -91.1 ms, Benjamini-Hochberg FDR-corrected p = 0.008). (E) Coefficients from the joint spectral model adding concurrent delta, theta, alpha and beta power changes. Only exponent change showed an association with reaction time slowing (β = -85.9 ms, FDR-corrected p = 0.006 across the five-term spectral family). In A to C, points are individual participants. In A to E, bands and whiskers are 95% confidence intervals. In D and E, significant terms are in blue and non-significant in grey. n = 222 (training set) in A to E. *p < 0.05, **p < 0.01, ***p < 0.001.

To test whether this effect was specific to the aperiodic exponent rather than to oscillatory features, we regressed the within-person change in mean reaction time on the concurrent change in the exponent and in delta, theta, alpha and beta band aperiodic-adjusted power, together with the baseline level of each spectral term, baseline age, baseline reaction time, sex and the follow-up interval. Of the five simultaneous spectral-change terms, only the exponent change predicted reaction time change after correction (β = -85.9 ms, 95% CI [-137.38, -34.45], FDR-corrected p = 0.006 across the five-term spectral family; adjusted R^2^ = 0.22) (**Fig. 3E**) (see **Supplementary Table 3**).

To confirm that this finding was robust to spectral-parameterization fitting ranges and regional electrode groups, we refitted the full coupling model across all six electrode groups, both eyes-open and eyes-closed conditions, and three spectral fitting ranges (1-35, 1-40, 1-45 Hz). The eyes-open exponent relationship was negative in every case and reached or approached significance across most electrode groups (**Supplementary Fig. 2A**). In the eyes-closed condition the effect was absent everywhere. It was never moderated by baseline age and did not depend on the follow-up interval, and baseline reaction time remained the only significant covariate (all other terms p > 0.05). Furthermore, although the exponent effect was regionally broad, it was not always the only spectral change to track slowing. Repeating the joint spectral model across the same condition, region and fitting-range grid, the change in the exponent was the most consistent and widely distributed spectral predictor of slowing under eyes-open but not eyes-closed recording. Some aperiodic-adjusted peak-power changes showed effects that were only locally over occipital electrodes (alpha and beta in eyes-open condition; delta, in eyes-closed condition) (**Supplementary Fig. 2B**).

### Simon task response times slow and ‘No-Go’ accuracy improves in neurotypical adults

We next characterized the longitudinal change in executive function, assayed by the Simon task across three measures: mean ‘Go’ response time, the Simon effect (incongruent minus congruent response time) indexing interference control, and ‘No Go’ accuracy indexing response inhibition. Within individuals, mean ‘Go’ response time increased from baseline to follow-up (mean change = 12.397 ms, 95% CI [4.49, 20.31]; permutation sign-flip test p = 0.002, paired t(221) = 3.09, d_z_ = 0.21), however the Simon effect did not change (mean change = 1.163 ms, 95% CI [-2.27, 4.60], permutation sign-flip test p = 0.50, d_z_ = 0.045). In contrast, ‘No Go’ accuracy improved (mean change = 0.029, 95% CI [0.01, 0.04]; permutation sign-flip test p < 0.001, paired t(221) = 3.68, d_z_ = 0.25) (**Fig. 4A-C**). Linear mixed-effects models of each measure (*measure ∼ session + age + sex + (1 | participant)*) confirmed these within-person changes and separated them from the between-person age gradient. Cross-sectionally, older baseline age was associated with slower response time (β = 2.227 ms per year, p < 0.001), a larger Simon effect (β = 0.448 ms per year, p < 0.001), and a modest reduction in no-go accuracy (β = -0.001 per year, p = 0.053) (**Fig. 4D-F**). Taken together, there is a clear dissociation across measures such that response times slowed both within individuals and across the cross-sectional age range. While the Simon effect carried a cross-sectional age gradient there was no within-person change, and yet ‘No-Go’ accuracy improved within individuals despite a marginal cross-sectional decline with age (**Fig. 4G-I**) (see **Supplementary Table 2**).

**Figure 4.**
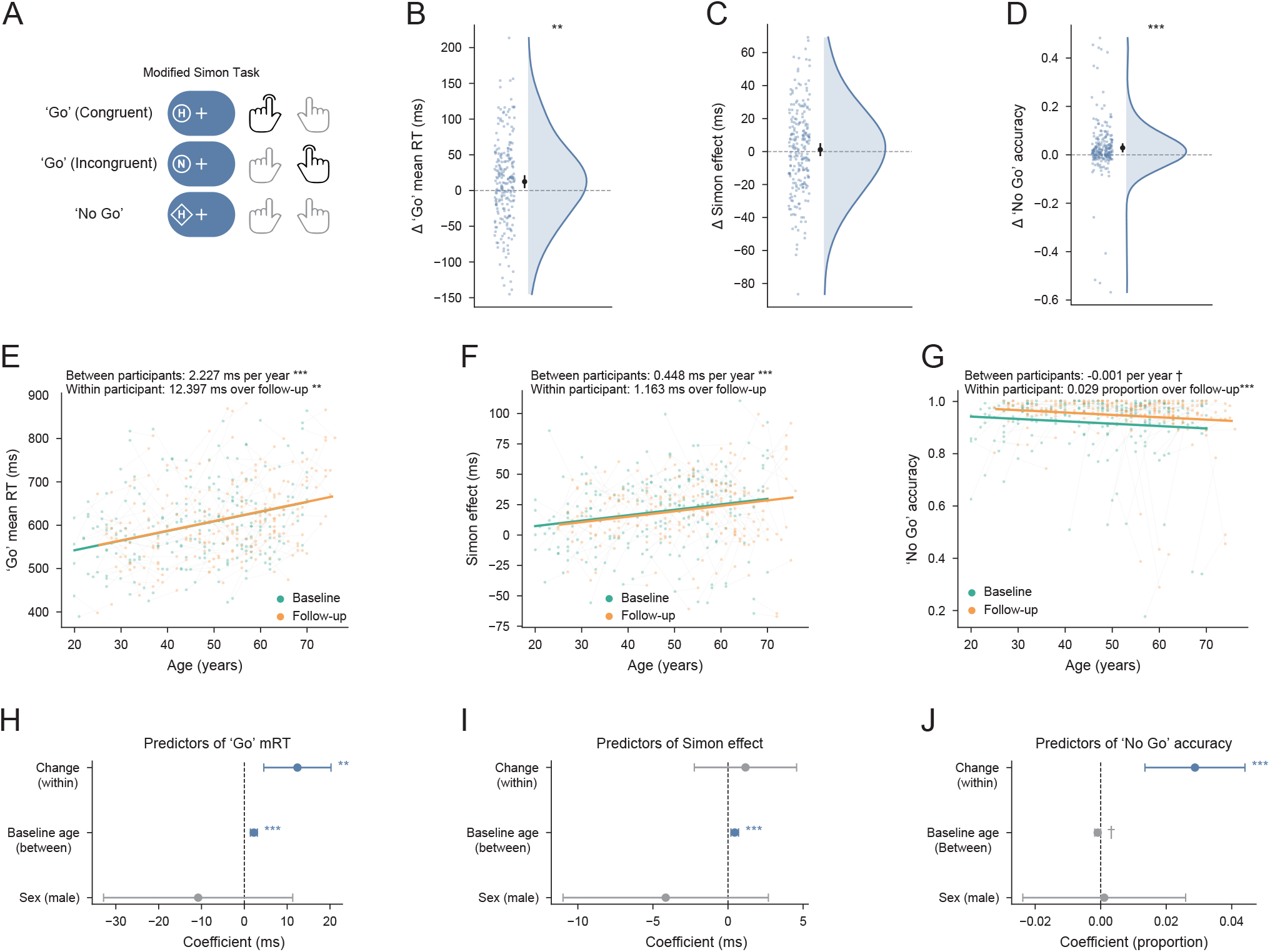
Response time in the modified Simon task slows and no-go accuracy improves within neurotypical adults. (A) Schematic of the modified Simon Task. On each trial a single letter target (‘H’ or ‘N’) appeared to the left or right of a central fixation cross. On ‘Go’ trials, indicated by a surrounding circle, participants responded with the left hand to ‘H’ and with the right hand to ‘N’, irrespective of the side of stimulus presentation. ‘Go’ trials were congruent when the stimulus appeared on the same side as the responding hand and incongruent when it appeared on the opposite side; the Simon effect was defined as the mean response time difference between incongruent and congruent ‘Go’ trials. On ‘No Go’ trials, indicated by a surrounding diamond, participants were instructed to withhold their response. (B-D) Within-person change (follow-up minus baseline) in (B) overall ‘Go’ response time, (C) the Simon effect and (D) ‘No Go’ accuracy. Clouds show the participant distribution and the point and whisker show the mean and its 95% confidence interval. Response time increased (mean = 12.397 ms, two-sided sign-flip permutation test, 10,000 permutations, p = 0.002) and ‘No Go’ accuracy improved (mean = 0.029, p < 0.001), whereas the Simon effect did not change (mean = 1.163 ms, p = 0.50). (E-G) Each measure against age at both sessions, baseline (teal) and follow-up (orange), with mixed-effects lines overlaid. In linear mixed-effects models, (E) older baseline age was associated with slower response time (β = 2.227 ms per year, p < 0.001), (F) a larger Simon effect (β = 0.448 ms per year, p < 0.001) and (G) marginally lower no-go accuracy (β = -0.001 per year, p = 0.053). Within participant effects are overlaid (in grey) as assessed in B-D. (H-J) Coefficients for each measure from a linear mixed-effects model (within-person change, baseline age, sex). Error bars denote 95% confidence intervals, significant terms are in blue and non-significant in grey. n = 222 (training set) in B to J. † < 0.07, **p < 0.01, ***p < 0.001.

We then confirmed that the two within-person changes were not artifacts of baseline-dependent change or session interval. For mean ‘Go’ response time, baseline performance predicted the magnitude of change (β = -0.213, 95% CI [-0.30, -0.13], p < 0.001) (**Supplementary Fig. 3A**), consistent with regression to the mean. Yet the slowing survived adjustment for baseline level (intercept at the mean baseline = 10.8 ms, 95% CI [1.05, 20.60], p = 0.03) and, unlike the cohort-wide PVT slowing, was concentrated in older participants, with this age relationship persisting after residualizing on baseline performance, sex and follow-up interval (β = 1.23 ms per year, 95% CI [0.64, 1.82], p < 0.001) (**Supplementary Fig. 3B**). ‘No Go’ accuracy likewise showed regression to the mean (β = -0.617, p < 0.001, adjusted R^2^ = 0.37) (**Supplementary Fig. 3C**) and survived baseline adjustment (intercept = 0.030, p < 0.001), but this improvement carried only a modest residual age relationship, such that older participants improved slightly less (β = -0.001 per year, 95% CI [-0.002, -.0000779], p = 0.034) (**Supplementary Fig. 3D**). Neither change depended on the follow-up time interval (both p > 0.14).

### The aperiodic exponent does not track concurrent Simon task measures

We next investigated whether the aperiodic exponent was correlated with Simon task performance, taking mean ‘Go’ response time as the executive-function analogue of PVT mean reaction time since it also slowed within individuals (**Fig. 4**). Across the pooled baseline and follow-up sessions, ‘Go’ response time did not correlate with the exponent (β = 16.7 ms per unit, 95% CI [-29.70, 63.04], p = 0.48) and, unlike the PVT, the slope relating the exponent to response time did not change at follow-up (*exponent x session*, β = -43.0 ms, 95% CI [-95.65, 9.70], p = 0.11) (**Fig. 5A**). Within individuals, exponent change was unrelated to change in mean ‘Go’ response time (r = -0.02, p = 0.78) (**Fig. 5B**), and baseline exponent likewise did not predict mean ‘Go’ response time change (r = -0.03, p = 0.69) (**Fig. 5C**). In a multivariable model that included the change and the baseline exponent simultaneously, together with baseline age, baseline response time, sex and the follow-up interval, the exponent effect was absent (*Δexponent*, β = -15.2 ms, 95% CI [-71.69, 41.21], FDR-corrected p = 0.86; adjusted R^2^ = 0.15) (**Fig. 5D**). The same model recovered the previously observed structure: ‘Go’ mean response time slowing was concentrated in older participants (*age*, β = 1.3 ms per year, 95% CI [0.68, 1.99], p < 0.001) and baseline ‘Go’ mean response time negatively related to its own change (β = -0.26, p < 0.001), consistent with regression to the mean. These results collectively confirm that the absent exponent effect was not an artifact of a model unable to recapitulate our prior observations. This null held across all six electrode groups, both recording conditions and three fitting ranges, with no surviving correction in any configuration (**Supplementary Fig. 4**). Applied to the remaining Simon measures, the same model was likewise null after correction. The exponent did not track the Simon effect (*Δexponent*, β = -3.1, FDR-corrected p = 0.88) (**Fig. 5E**) nor ‘No Go’ accuracy (*Δexponent*, β = 0.084, FDR-corrected p = 0.70) (**Fig. 5F**). Thus, in contrast to the PVT, the within-person exponent change was uncoupled from every Simon measure.

**Figure 5.**
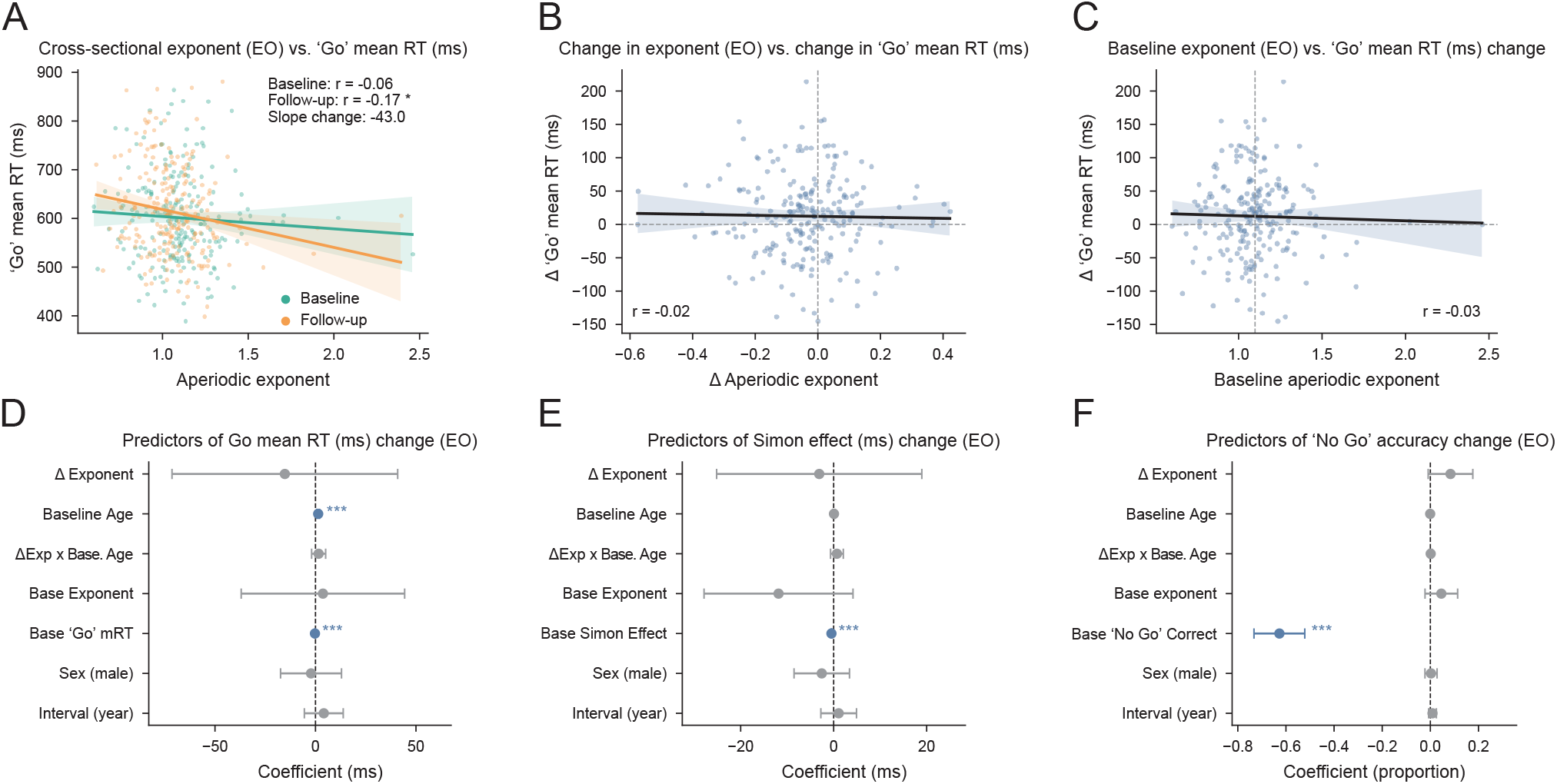
The aperiodic exponent does not track concurrent Simon measures. (A) ‘Go’ response time against the aperiodic exponent across pooled sessions at baseline (teal) and follow-up (orange). Lines and shading show the fit and 95% confidence interval from cluster-robust ordinary least-squares regression. The exponent did not relate to response time (β = 16.67, p = 0.48) overall and the slope did not change at follow-up (exponent x session β = -43 ms, p = 0.11). (B) Within-person Go response-time change against exponent change did not show a relationship (Pearson r = -0.02, p = 0.78). (C) Response-time change against the baseline exponent also did not show a relationship (Pearson r = -0.03, p = 0.69). (D) Coefficients from the full coupling model of ‘Go’ response-time change (ordinary least-squares with covariates: exponent change, baseline age, their interaction, baseline exponent, baseline reaction time, sex, follow-up interval). Exponent change was not associated with slowing (β = -15.2 ms, Benjamini-Hochberg FDR-corrected p = 0.86), though the model recovered age-related slowing (β = 1.3 ms per year, p < 0.001) and regression to the mean in baseline response time (β = -0.26 ms per year, p < 0.001). (E) The same coupling model for Simon effect change showed no coupling with the exponent (β = -3.1, FDR-corrected p = 0.88), with baseline Simon effect the only significant term (β = -0.479, p < 0.001). (F) The same coupling model for ‘No Go’ accuracy change also showed no coupling with the exponent (β = 0.084, FDR-corrected p = 0.70). Points in A to C are individual participants and the shaded band is the 95% confidence interval of the fit. In D to F, points are model coefficients and whiskers are 95% confidence intervals, with significant terms in blue and non-significant in grey. n = 222 (training set) in A to F. ***p < 0.001.

### Aperiodic exponent decline carries a stable, state- and domain-specific predictive signal for attentional slowing

The mixed-effects model results that we previously observed (**Fig. 3**) were estimated in a single sample, so we next asked whether it was stable, state-specific, and behaviorally specific. We thus refitted the model across 200 age-stratified 80/20 cross-validation splits of the training set (n = 222; n = 220 for post-task models, two participants each lacking a post-task eyes-open recording at one session). We used these repeated within-training splits to assess stability, reserving the frozen test set (n = 54) as the single independent generalization test.

Using pre-task data, the coefficient relating exponent change to PVT mean reaction time change was negative in every resample (i.e., a steeper exponent decline predicted greater slowing; median -93.46 ms per unit), and its sign agreed between the model-fitting and held-out partitions of each split in 93% of resamples (single-partition training estimate β = -91.11 ms, p < 0.001) (**Fig. 6A**) (see **Supplementary Table 5**). The post-task coefficient, in contrast, was centered near zero with an unstable sign (median -2.08 ms, negative in 56% of resamples, partition concordance 16%), indicating that the stable relationship present in pre-task EEG was absent in post-task EEG. To further explore whether this was due to state-specificity (e.g., cognitive or attentional fatigue over the entirety of the experiment), we tested the held-out prediction of the PVT change and found that both models predicted above zero in the majority of resamples (pre-task median held-out R^2^ = 0.167, root-mean-squared error = 51.1 ms, positive in 84% of resamples; post-task median held-out R2 = 0.122, root-mean-squared error = 52.0 ms, positive in 74%) (**Fig. 6B**). However, both were largely driven by baseline reaction time (see **Fig. 3D**), suggesting that regression to the mean inflated this marginal R^2^ and thus obscured the exponent’s specific contribution.

**Figure 6.**
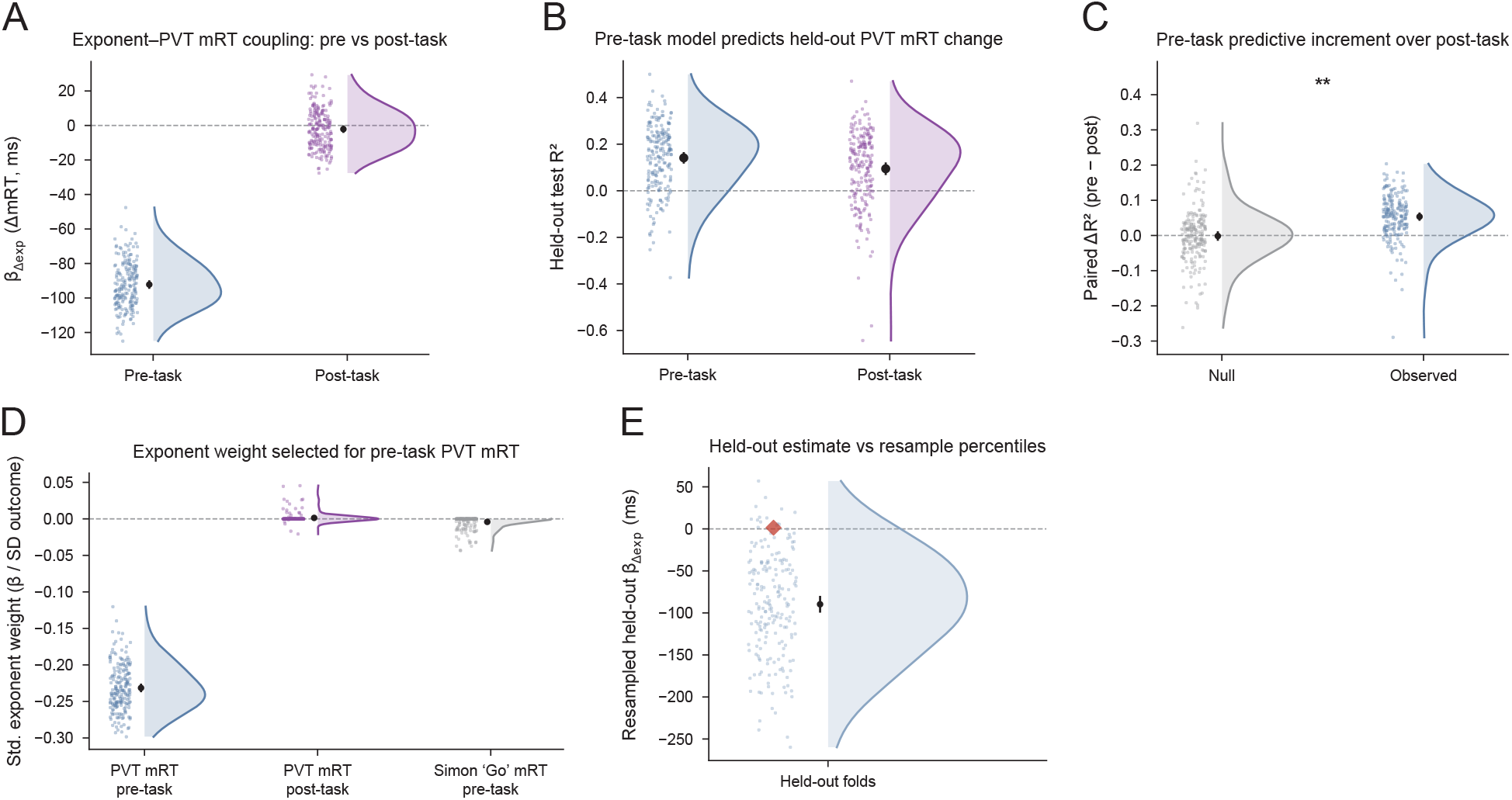
The aperiodic exponent and PVT slowing coupling is stable across training resamples, state-specific and domain-specific. Out-of-sample prediction of within-person PVT reaction time change across 200 age-stratified 80/20 resamples of the training set, fitted separately to pre-task (blue) and post-task (purple) EEG. Clouds show the distribution across resamples, and the point and whisker show the mean across resamples and its 95% confidence interval. (A) Ordinary least-squares coefficient relating exponent change to PVT reaction time change. The pre-task training-partition coefficient was negative in every resample (median = -93.46 ms) with 93% sign concordance between the independently fitted training and held-out partitions, whereas the post-task coefficient was near zero with an unstable sign (median = -2.08 ms, negative in 56.5% of resamples, 16% concordance). (B) Held-out R^²^ of each model on the held-out partition of each resample (pre-task median R^²^ = 0.167, RMSE = 51.1 ms, positive in 84% of resamples; post-task median R^²^ = 0.122, RMSE = 52.0 ms, positive in 74%). Both arms are largely driven by baseline reaction time (Fig. 3D). The paired difference in C isolates the pre-task contribution. (C) Paired difference in held-out R² (pre-task minus post-task) on the same participants (blue) beside a null obtained by permuting the outcome within each resample’s training partition (grey). The mean paired increment exceeded a full label-shuffle null in a separate two-sided permutation test (observed mean ΔR^²^ = 0.052; null mean = -0.001, SD = 0.018; 1,000 global permutations x 100 folds, p = 0.008); asterisks denote that permutation test. Across the 200 resamples the increment had a median of 0.058 (mean 0.053) and a 2.5th-97.5th percentile spread of [-0.078, 0.162]. (D) Standardized elastic-net exponent weight (coefficient divided by the within-fold outcome standard deviation) for PVT reaction time pre-task (blue), PVT reaction time post-task (purple) and Simon ‘Go’ response time pre-task (grey). The pre-task PVT weight was consistently negative and survived the L1 penalty in 100% of resamples (median = -0.235). The post-task PVT weight and the pre-task Simon weight were negligible (median approximately 0, retained in 12% and 28% of resamples, respectively). (E) The coupling coefficient refitted within the frozen 54-person held-out set (red diamond) against the distribution of held-out-partition coefficients from the training-set resamples (blue). The frozen coefficient was positive and non-significant (β = 1.416, p = 0.983), falling at the 93rd percentile of the resample distribution, and the out-of-sample R^²^ of the frozen training model applied to the held-out set was negative (-0.181, worse than predicting the held-out mean), at the 2.5th percentile. Generalization is therefore supported by coefficient stability within the training sample rather than by the independent held-out test. n = 222 (training, resampled) for pre-task panels; n = 220 for post-task panels and for all paired pre-minus-post comparisons, two participants each lacking a post-task eyes-open recording at one session; n = 54 (frozen held-out set). **p < 0.01.

To remove that shared component, we took the paired difference in held-out R^2^ between the pre-task and post-task models fitted on the same 220 participants. This pre-minus-post increment was positive (median across resamples 0.058, mean 0.053, 2.5th-97.5th percentile spread [-0.078, 0.162]), and its mean exceeded a full label-shuffle null (observed mean ΔR^2^ = 0.052; null mean = -0.001, SD = 0.018; two-sided permutation test, 1,000 global permutations x 100 folds, p = 0.008) (**Fig. 6C**), indicating a modest, reliable predictive advantage of pre-task over post-task EEG for the PVT change relationship. We then asked whether state-specificity also manifested at the level of feature use. We fitted a penalized (elastic-net) model and expressed the exponent weight on a common scale across sessions (standardized by the outcome’s standard deviation). The pre-task exponent carried a consistent negative weight that survived the L1 penalty in 100% of resamples (median standardized weight -0.235), whereas the post-task exponent weight was approximately zero and retained in only 12% of resamples (**Fig. 6D**). Placing the exponent weight on a common scale, the pre-task exponent predicting change in Simon ‘Go’ response time was negligible and rarely selected (median standardized weight approximately 0, retained in only 28% of resamples), in contrast to the pre-task PVT weight (median -0.235), which was retained in every resample, indicating a domain dissociation.

We then evaluated the *a priori* held-out set (n = 54) as the single independent test of generalizability (**Fig. 1A**). Refitting the coupling model within the held-out set returned a non-significant, positive coefficient (β = 1.416 ms, p = 0.98), and applying the frozen training model to predict held-out PVT change gave a negative out-of-sample R^2^ (-0.181), indicating a prediction worse than the held-out sample mean. These fell at the 93rd and 2.5th percentiles, respectively, of the resample distribution **(Fig. 6E)**. The coefficient being in the upper tail of the distribution (where the held-out coefficient flips sign) combined with an R^2^ near the bottom suggests an underpowered draw, since cross-validated estimates carry large sampling variance at small sample sizes^52^. Thus, our one fully independent test did not confirm out-of-sample prediction, and we therefore present generalization as supported by coefficient stability within the training sample only.

Repeating the full analysis with folds stratified jointly on age and baseline mean response time (in place of age alone) led to similar findings: the pre-task coefficient remained negative in 100% of resamples (median -91.88 ms, concordance 100%), the paired predictive increment was again significant yet modest (observed mean ΔR^2^ = 0.056, two-sided permutation p = 0.014), and the standardized exponent weights reproduced the same state and domain dissociations (PVT pre-task median standardized weight = -0.232, retained in 100% of resamples; PVT post-task median standardized weight approximately 0, retained in 12.5% of resamples; Simon pre-task median standardized weight approximately 0, retained in 22.5% of resamples) (**Supplementary Fig. 5**).

## Discussion

Our analyses asked whether within-person changes in the task-free EEG aperiodic exponent track concurrent cognitive-behavioral changes across aging, and whether any such coupling is generalizable across cognitive-behavioral domains. We found that the aperiodic exponent declined within individuals over an approximately five-year follow-up interval, separably from the age at baseline and from concurrent changes in aperiodic adjusted oscillatory-power, and robustly across spectral parameterization fitting ranges and electrode regions. In parallel, PVT mean reaction time slowed over the same interval, reflecting age-related decline in sustained attention and processing speed, while reaction-time variability and accuracy did not change. Our central finding is that the within-person exponent decline tracked concurrent PVT slowing after adjusting for multiple covariates, including baseline age and performance, and was specific to the aperiodic exponent rather than to oscillatory features. By contrast, no exponent relationship emerged for any Simon-task measure despite behavioral changes in that domain, including ‘Go’ response time slowing and ‘No-Go’ accuracy improvements, suggesting a cognitive-behavioral domain dissociation. Our predictive modeling analyses showed that the exponent decline and PVT slowing coupling was stable across resamples, was state-specific (pre-task over post-task EEG), and was domain-specific (PVT over Simon task performance). Overall, our findings suggest that the age-related aperiodic exponent decline indexes attentional and processing-speed performance rather than general cognitive-behavioral aging, even within a time interval as short as five years.

Prior cross-sectional work established that the slope of the EEG power spectrum in log-log space ‘flattens’ with neurotypical aging (i.e., the aperiodic exponent decreases)^10,22,35^. An independent study drawing on the same cohort we analyzed (the Dortmund Vital Study^46,47^) similarly showed that the exponent declines over an average of five years in neurotypical adults^24^. Here, we extend these findings as the first to show that within-person change in the aperiodic exponent tracks concurrent cognitive-behavioral change within the same individuals, and report that this coupling may be dissociated from cross-sectional age relationships. For instance, the between-person age gradient sometimes ran opposite to the within-person change (e.g., on PVT mean RT, older baseline age was associated with faster responding, yet reaction time nonetheless slowed within individuals across the interval) and sometimes existed without it (the Simon effect carried an age gradient but no longitudinal change). This dissociation aligns with findings from the largest cross-sectional study of aperiodic activity and cognitive aging to date, in which the exponent declined with age across more than 1700 older adults^53^. That study found no reliable association with cognition, including a processing-speed factor indexed by the Stroop test^53^. The coupling we report here relies on within-person changes rather than the static exponent effect observed in cross-sectional study designs. Similarly, the baseline exponent did not predict PVT change in our data (**Fig. 3C**). These dissociations caution against inferring within-person aging trajectories from cross-sectional age effects in studies of aperiodic neural activity and cognition.

The within-person exponent decline tracked PVT slowing but not any change in Simon-task measures, including ‘Go’ response time, which, while it slowed longitudinally, remained uncoupled. This suggests that the exponent tracks the specific attentional-speed demands of the PVT rather than general RT slowing. But executive function also recruits information processing, so one possibility, which our work does not directly test, is that the aperiodic exponent tracks the speed or efficiency of information processing shared across cognitive-behavioral tasks, rather than interference control itself. For example, a ‘flatter’ spectral slope in older adults is linked to less consistent stimulus-evoked responses in a visual attention task^26^ and to sustained attention specifically^29^. These studies, combined with the absence of exponent coupling with the Simon effect (**Fig. 5E**) and the faint, same-direction Simon ‘Go’ reaction-time signal in our predictive analysis (**Fig. 6D**), fit the idea that the aperiodic exponent indexes processing-speed demands shared across cognitive-behavioral tasks. Furthermore, cross-sectional work has shown that the aperiodic exponent relates to some cognitive domains more than others^37,40^, so a domain-specific coupling may not be atypical.

Extending beyond domain specificity toward state specificity, the coupling between the aperiodic exponent and PVT reaction time was carried by pre-task, but not post-task, EEG (**Fig. 6A, D**) and was present in the eyes-open, but not eyes-closed, condition (**Supplementary Fig. 2A**). It may be that EEG collected before any task is reflective of a rested baseline that is more predictive of subsequent behavior, whereas EEG after the task could be contaminated by task-induced fatigue and general shifts in arousal^38,39^. The pre-minus-post-task predictive increment reached significance against a full label-shuffle null (observed mean ΔR^2^ = 0.052, p = 0.008) (**Fig. 6C**), although it was modest in absolute magnitude. This was further supported by a coupling coefficient that was sign-stable in 100% of resamples, and an exponent weight that survived the L1 penalty in 100% of pre-task versus 12% of post-task resamples, consistent with state-specific exponent-PVT mRT coupling.

A key limitation in interpreting our results stems from the observational and correlational nature of our findings. We therefore cannot establish that exponent decline causes attentional slowing, as both may arise from upstream disruptions found in healthy aging, such as neurodegeneration or vascular changes. Aperiodic activity is also a complex measure with many neural contributors^18^, as well as several non-neural sources as well^54^. Most notably, cardiac activity can survive ICA-based correction and has been shown to influence aperiodic exponent estimates^55^. However, such non-neuronal physiological confounds should track behavior irrespective of cognitive domain, recording state, or eyes condition; our observed coupling was specific to the PVT, to pre-task EEG, and to the eyes-open condition, suggesting that the coupling is not reducible to non-neural contributions to the aperiodic exponent. With regards to our *a priori* held-out test set, the refitted coefficient was positive and non-significant (β = 1.416, p = 0.98), sitting at the 93rd percentile of the resample distribution, while the out-of-sample R^2^ was negative (-0.181) at the 2.5th percentile (**Fig. 6E**), suggesting an atypical, low-powered draw rather than a clear null, though generalization to the independent set was not confirmed. Our generalization claims therefore rest on the resampling distribution and feature-weight analyses.

An additional challenge in interpreting our findings is that the healthy-volunteer cohort produced a counterintuitive between-person age effect (older participants faster at baseline on RT) (**Fig. 2C, D**), which may reflect an inherent recruitment bias toward healthier and more cognitively able older adults in studies of this kind, limiting generalization to clinical or at-risk populations. Furthermore, education has been shown to moderate the cross-sectional relationship between the aperiodic exponent and cognition^36^. Education may be more likely a moderator of the rate of change than a confound of the coupling itself, though with only two timepoints we cannot characterize the trajectory’s shape, non-linearity, or heterogeneity in the rate of change across our measures. Finally, our reported effect sizes were modest (d_z_ values of approximately 0.2 to 0.3) and the exponent’s incremental predictive contribution beyond baseline covariates was small in absolute terms. With only two cognitive-behavioral domains tested, there remains a clear need to delineate what the aperiodic exponent does and does not track and whether any changes are advantageous to daily life tasks.

A non-invasive, low-cost, scalable measure of within-person neural change with decent test-retest reliability^24^ is well suited to screening and monitoring healthy and compromised aging, particularly where repeated visits are feasible. Our study shows that within-person exponent changes track concurrent cognitive-behavioral changes, positioning the aperiodic exponent as a longitudinal, within-person neural biomarker of attentional slowing rather than a cross-sectional correlate. The exponent is widely interpreted as an index of cortical excitation-inhibition balance, such that a flatter spectrum reflects a shift toward excess excitation and reduced inhibition with age^10,15^. While we do not test that framework here, we provide a foundation for pairing aperiodic exponent measures with more direct indices of synaptic and neurotransmitter tone, such as its reported relationship to glutamatergic signaling^32^, to track changes in attentional performance in model organisms. Beyond mechanism, future work should extend these findings to clinical and at-risk cohorts such as mild cognitive impairment and neurodegenerative disorders, use a denser longitudinal sampling to model individual trajectories, broaden the cognitive battery to map the boundaries of aperiodic neural activity coupling, and combine the exponent with complementary, multimodal biomarkers such as white-matter microstructure. The urgent need for objective, scalable, within-person neural markers of cognitive change in aging populations positions the longitudinal aperiodic exponent as a candidate marker that, as our findings suggest, is specific to attentional aging.

## Methods

### Participants and study design

We analyzed data from a finalized count of 276 neurotypical adults from the Dortmund Vital Study^46,47^, an ongoing longitudinal cohort study of cognitive aging across adults aged 20 to 70 years conducted at the Leibniz Research Centre for Working Environment and Human Factors at the Technical University of Dortmund, Germany (IfADo; ClinicalTrials.gov NCT05155397). Each analyzed participant had complete task-free EEG and cognitive-behavioral assessments at two timepoints approximately five years apart (5.3 ± 0.8 years; mean ± SD). Participants were recruited from local companies and public institutions and through public advertisements.Exclusion criteria comprised a history of severe disease, namely neurological disease (such as dementia, Parkinson’s disease or stroke), cardiovascular, oncological and eye disease, psychiatric and affective disorders, head injury, head surgery or head implants, the use of psychotropic drugs and neuroleptics, and limited physical fitness or mobility. Common medications such as blood thinners, hormones, antihypertensives and cholesterol-lowering agents were not grounds for exclusion. Participants reported being healthy and free of medication that could affect attention during the experimental sessions. All participants gave written informed consent and the study conformed to the Declaration of Helsinki with approval by the local Ethics Committee of the Leibniz Research Centre for Working Environment and Human Factors, Dortmund (approval A93-1, and A93-3 for the follow-up). The present sample additionally draws on follow-up data beyond the original release (see Data availability). Participant’s age at second session was not exact, but rather reconstructed from the recording years in the BIDS session files as the baseline age plus the elapsed interval.

### Cognitive-behavioral assessments

At each session participants completed a standardized battery of five cognitive tasks; the present analyses used two of these, indexing distinct domains: Sustained attention and processing speed were measured with a ten-minute psychomotor vigilance task (PVT)^48,49^ adapted to present the target (“+”) at one of four random interstimulus intervals (2, 3, 5 or 8 s), with the screen blank during the interval. Participants responded to each target onset with a button press as quickly as possible. We computed mean reaction time, reaction-time variability and the proportion of correct responses, each averaged across the five consecutive two-minute blocks of the ten-minute task. Executive function, interference control, and response inhibition were measured with a modified version of the Simon task comprising intermixed ‘Go’ and ‘No-Go’ trials^50,51^ On each trial a target appeared to the left or right of fixation, and participants responded with a left- or right-hand keypress according to the non-spatial target feature (letter “H” or “N”) while the task-irrelevant target location varied independently. ‘No-Go’ trials, signaled by a surrounding diamond, required the response to be withheld. Among ‘Go’ trials, signaled by a surrounding circle, we distinguished congruent trials (target location and response side on the same side) from incongruent trials (opposite sides) and defined the Simon effect as the difference in mean response time between incongruent and congruent ‘Go’ trials. We used mean ‘Go’ response time across all ‘Go’ trials, and ‘No-Go’ accuracy (the proportion of ‘No-Go’ trials on which the response was correctly withheld) as an index of response inhibition.

### EEG data acquisition

Task-free scalp EEG was recorded with a 64-channel elastic cap, electrodes connected with saline solution and arranged according to the international 10-20 system, using a BrainVision BrainAmp DC amplifier and BrainVision Recorder software (Brain Products GmbH) with channel ‘FCz’ as the online reference. Signals were digitized at 1000 Hz with a 250-Hz online low-pass filter and no online high-pass filter, and electrode impedances were kept below 10 kOhm. At each session, task-free EEG was acquired for three minutes with eyes closed and three minutes with eyes open, always in that order, and this pair of recordings was obtained both before (pre) and after (post) the approximately two-hour cognitive-behavioral task battery, yielding four resting recordings per session roughly two hours apart. All testing was conducted in the morning at a fixed time across participants to standardize wakefulness, and the pre-task and post-task recordings allowed us to test the state specificity of any brain-behavior coupling.

### EEG preprocessing

Preprocessing was performed in MNE-Python^56^ (see the Software section for all package versions used in our analyses). Continuous recordings were assigned the 64-channel 10-20 montage. Line noise was attenuated with a spectrum-fit notch filter at 50 Hz and its first harmonic (100 Hz), and the data were band-pass filtered between 1 and 100 Hz with a zero-phase FIR filter (Hamming window) before re-referencing from the ‘FCz’ online reference to the common average. Ocular, muscular and other non-neural artefacts were removed with independent component analysis (extended Infomax, 20 components), with the decomposition fitted under a peak-to-peak amplitude ‘global’ rejection threshold estimated by AutoReject^57^ so that high-amplitude segments did not dominate the solution. Components were classified with ICLabel^58^, and all components not labeled ‘brain’ or ‘other’ were discarded before back-projection. The post-processed data were segmented into non-overlapping ten-second epochs, and residual bad epochs were rejected or repaired with AutoReject (via calculation of local high-amplitude rejection thresholds). Finally, a 100-ms Hamming taper was applied to the leading and trailing edges of each epoch to limit spectral leakage that may occur when non-consecutive post-processed epochs were appended.

### Spectral parameterization

Power spectral densities were estimated from the artifact-corrected epochs using Welch’s method (one-second Hamming window, 50% overlap, two-second FFT length, median across windows) over 1 to 80 Hz, and the resulting per-epoch spectra were combined across epochs by their median to give a single spectrum per channel that was then parameterized with the *specparam* algorithm (formerly FOOOF)^9^, fitted over 1 to 45 Hz in the primary analysis, with peak width limits of 2 to 12 Hz, no limit on the number of peaks, a minimum peak height of 0 and a peak threshold of 2 standard deviation. For every channel we fitted two aperiodic models, one assuming a single power law (‘fixed’ mode) and one incorporating a spectral knee (‘knee’ mode). To confirm that the choice of fit range did not drive results, the full pipeline was repeated with upper fit bounds of 35 and 40 Hz as a sensitivity analysis. While the aperiodic exponent was the primary feature of interest, we additionally computed the aperiodic-adjusted peak power within delta (1 to 4 Hz), theta (4 to 7 Hz), alpha (7 to 13 Hz) and beta (13 to 24 Hz) bands, specifically the height of the largest spectral peak above the aperiodic fit within each band.

Because a spectral ‘knee’ may or may not be visually present in each power spectra, we selected the *specparam* aperiodic mode per participant and per task rather than imposing this mode on the whole sample. Therefore, for each participant-task pair we compared the model outputs from individually fitted ‘fixed’ and ‘knee’ modes using the adjusted R^2^ of the fit over a low-frequency subrange (1 to 20 Hz), averaged across that participant’s baseline recordings. The ‘knee’ model was adopted only when it improved adjusted R^2^ over the ‘fixed’ model and its estimated ‘knee’ frequency fell within the fitted range, suggesting the presence of a biophysically plausible ‘knee’, otherwise the ‘fixed’ model was retained. The selected mode was then applied to all that participant-task pair’s recordings across both sessions, and these mode-validated outputs, such as the aperiodic exponent, were used in all primary analyses.

Model fits were also subjected to a channel-level quality gate before any averaging: channels whose fit explained less than 90% of the spectral variance (R^2^ below 0.9) were removed from the average. Summaries were formed for five scalp EEG electrode regions (frontal: ‘Fp1’, ‘Fp2’, ‘F3’, ‘AF3’, ‘F5’, ‘F1’, ‘FC3’, ‘F4’, ‘AF4’, ‘F6’, ‘F2’, ‘FC4’, ‘Fz’; temporal: ‘T7’, ‘FT7’, ‘TP7’, ‘T8’, ‘FT8’, ‘TP8’; central: ‘C3’, ‘C5’, ‘C1’, ‘CP3’, ‘C4’, ‘C2’, ‘C6’, ‘CP4’, ‘Cz’; parietal: ‘P3’, ‘P5’, ‘P1’, ‘P7’, ‘Pz’, ‘P4’, ‘P6’, ‘P2’, ‘P8’; occipital: ‘O1’, ‘O2’, ‘Oz’, ‘PO7’, ‘PO3’, ‘POz’, ‘PO4’, ‘PO8’), each requiring at least two surviving channels to be computed, and together with the whole-scalp average these constituted the six electrode groupings used in the regional robustness analyses. A participant was flagged for exclusion if fewer than 32 channels survived the gate in each recording (n=17 participants, not included in the final count). Unless otherwise stated, analyses used the whole-scalp average exponent across surviving channels for a total of 276 participants.

### Statistical analyses

#### Held-out validation split

To provide an unbiased test of predictive generalization, we defined a single train/test split *a priori* and froze it for all downstream analyses. Participants with complete pre-task EEG recordings in both conditions at both sessions, complete behavioral data at both sessions, and no recording excluded by the R^2^ gate were eligible. We allocated 20% of eligible participants to a held-out test set (n = 54) and the remaining 80% to a training set (n = 222), stratifying the allocation by baseline age band (under 30, 30 to 50, 50 to 70 and 70 or over years) so that the age distribution was preserved across sets. The split was generated once with a fixed random seed and stored as a fixed reference and no subsequent analysis altered it. The two sets were compared on baseline age with a Mann-Whitney U test and on sex with a chi-squared test and did not differ on either. All analyses, unless stated otherwise, used the training set (n = 222).

#### Longitudinal and cross-sectional effects

Aperiodic exponent and behavioral measures were tested to see whether they changed across the follow-up. Within-person changes were tested with two-sided paired t-tests and corroborated with sign-flip permutation tests (see Permutation tests). Effect sizes were reported as Cohen’s d_z_, with all differences signed as follow-up minus baseline. To confirm that any within-person change was not an artefact of baseline-dependent change or of the variable follow-up interval, we regressed each change score on its own baseline level and the follow-up interval and inspected the intercept at the mean baseline. We then revisualized the change on baseline level, sex and interval and tested its dependence on baseline age. To confirm that the exponent decline was not a by-product of concurrent oscillatory power changes, we regressed the within-person exponent change on the concurrent within-person changes in delta, theta, alpha and beta aperiodic-adjusted peak power, with baseline age and sex.

To separate the between-person age gradient from within-person change, we fitted linear mixed-effects models of each EEG and behavioral measure with a fixed effect of session, baseline age and sex and a by-participant random intercept (*measure ∼ session + age + sex + (1 | participant)*). The session term estimated the within-person change across the follow-up and the age term estimated the cross-sectional gradient. Raw coefficients (β) are reported in natural units with 95% confidence intervals and exact p-values, unless stated otherwise.

#### Brain-behavior coupling and state specificity

Our central analysis asked whether within-person change in EEG measures tracked concurrent change in each cognitive-behavioral measure. Because the exponent declined comparably under eyes-open and eyes-closed conditions, we focused our primary analyses on the eyes-open condition unless stated otherwise. Across the pooled baseline and follow-up sessions we first related the measure to the exponent with ordinary least squares using cluster-robust standard errors (clustered by participant), including an exponent-by-session interaction to test whether the cross-sessional relationship changed over time. The primary test then used within-person change scores, computed as follow-up minus baseline for every measure: we regressed cognitive change on exponent change, adjusting for baseline cognitive performance, baseline exponent, baseline age, sex and the per-person follow-up interval. Baseline age was mean centered within the analysis sample (n=222) rather than across the full cohort (n=276) until predictive analyses were performed. Coefficients are reported in their natural units, for example milliseconds of reaction-time change per unit change in the exponent, with 95% confidence intervals and exact p-values unless p < 0.001.

To test whether coupling was specific to the aperiodic exponent rather than to oscillatory features, we additionally regressed each cognitive-behavioral change on the concurrent change in the exponent together with the concurrent changes in delta, theta, alpha and beta aperiodic-adjusted peak power, the baseline level of each spectral term and the same covariates and controlled the false discovery rate across this five-term spectral family. For this model the concurrent band-power changes were each divided by their standard deviation (expressed in per-standard-deviation units) so that their coefficients would be comparable as effect sizes, while the outcome and the age and sex terms remained in raw units. All change-score and coupling models were refitted across the three spectral fitting ranges (1-35, 1-40 and 1-45 Hz), the six electrode groupings and both recording conditions for a sensitivity analysis.

To test whether coupling depended on recording condition (eyes-open and eyes-closed) we estimated it separately with linear mixed-effects models and tested the between-condition difference with a single model carrying a condition-by-exponent interaction, reporting the within-condition estimates from the separate per-condition models. State specificity was examined in the same way by contrasting the pre-task and post-task sessions.

#### Multiple-comparison correction and permutation tests

Where a family of related tests was performed, we controlled the false discovery rate (q < 0.05) with the Benjamini-Hochberg procedure, defining the membership of each family explicitly in the text. Effects surviving only an uncorrected threshold are not reported as significant (p > 0.05). We corroborated all within-person changes and coupling effects with non-parametric permutation tests. For each test the differences within-person were randomly multiplied by plus or minus one across 10,000 permutations and the one-sample t-statistic was recomputed. The two-sided p-value was thus the proportion of permuted statistics whose magnitude met or exceeded the initial observation. For the exponent and behavioral coupling models, the ΔExponent term was FDR-corrected across the family of six electrode regions x three task measures (18 tests), separately within the PVT and Simon analyses, whereas the joint spectral model was corrected across its five-term Δ-spectral family.

#### Predictive modelling and held-out validation

To test whether the within-person exponent decline that tracked PVT slowing reflected a stable, behaviorally specific signal rather than a single-sample association, we refitted the model across 200 age-stratified 80/20 resamples of the training set (n = 222), using the eyes-open, pre-task recordings unless stated otherwise. Two training participants were each missing a post-task eyes-open recording at one of the two sessions, so all post-task models, and the paired pre-versus-post comparisons that require both recordings from the same participant, were fitted to the 220 participants with complete pre- and post-task data. The held-out test set was complete for both recordings (n = 54). Each resample fitted the model on an 80% partition and evaluated it on the held-out 20%, so these repeated within-training splits index the stability of the coupling rather than independent generalization, for which we reserved the *a priori* held-out set (n = 54).

Every model in this framework used the brain-behavior coupling predictor set, namely the change in the exponent and its interaction with baseline age, the baseline exponent, baseline reaction time, baseline age, sex and the per-person follow-up interval, so that the resampled coefficient would be directly comparable with the primary coupling estimate. Within each resample the coupling coefficient was estimated separately in the training and held-out partitions, and we reported their sign concordance. To isolate the contribution of the exponent from the shared, baseline-driven prediction that arises through regression to the mean, we computed the paired difference in held-out R^2^ between the pre-task and post-task models on identical participants (the pre-minus-post increment) and tested it against a two-sided full label-shuffle null (1,000 permutations across 100 folds), the statistic being the mean ΔR^2^ across the 100 folds. To compare feature use across cognitive-behavioral domains on a common scale, we fitted a penalized (ElasticNet) model whose penalty was tuned by five-fold cross-validation on the training data (the L1 ratio searched over 0.1 to 1.0), with predictors standardized within each fold, and expressed the exponent weight after dividing by the within-fold outcome standard deviation, recording how often the weight survived the L1 penalty across resamples. As a confirmatory test on the *a priori* held-out set (n = 54), we refit the coupling model within the held-out set to obtain its coupling coefficient and separately applied the frozen training model to predict held-out reaction-time change to obtain an out-of-sample R^2^, both of which we report together with the percentile at which they fell in the training-resample distribution.

### Software

All preprocessing, spectral parameterization, statistical analysis and visualization were carried out in Python (3.9.19), using MNE-Python^56^ (1.8.0) for EEG handling and preprocessing, mne-icalabel^58^ (0.7.0) for ICLabel component classification, AutoReject^57^ (0.4.3) for epoch rejection and threshold estimation, *specparam*^9^ (2.0.0rc6) for spectral parameterization, statsmodels^59^ (0.14.4) for the regression and mixed-effects models, scikit-learn (1.4.2) for the ElasticNet models, h5py (3.11.0) for reading the behavioral data, and NumPy^60^ (1.26.4), pandas (1.5.3), SciPy (1.13.1), Matplotlib^61^ (3.9.0) and seaborn^62^ (0.13.2) for numerical work and plotting.

## Data availability

A subset of the EEG data analyzed in this study (n=208 participants, baseline release) is available at OpenNeuro (dataset accession number: ds005385, 10.18112/openneuro.ds005385.v1.0.2). The most up-to-date raw EEG and behavioral data (including the n = 276 used in this study after processing) is available upon reasonable request to Drs. Getzmann and Wascher.

## Code availability

All code used to recreate figures is publicly available on GitHub: https://github.com/voytekresearch/Adult_EEG_Dortmund.

## Acknowledgements

We’d like to thank Dillan Cellier and Dr. Maribel Patiño for their thoughtful discussion.

## Author Contributions

C.C.: Conceptualization, Formal analysis, Investigation, Methodology, Visualization, Writing - original draft, Writing - review and editing.

A.R.: Formal analysis, Writing - review and editing. L.K.: Formal analysis, Writing - review and editing

S.G.: Investigation, Methodology and Writing - review and editing S.A.: Investigation, Methodology and Writing - review and editing E.W.: Investigation, Methodology and Writing - review and editing

B.V.: Supervision, Project administration, Funding acquisition, and Writing - review and editing.

---

**Supplementary Figure 1.**
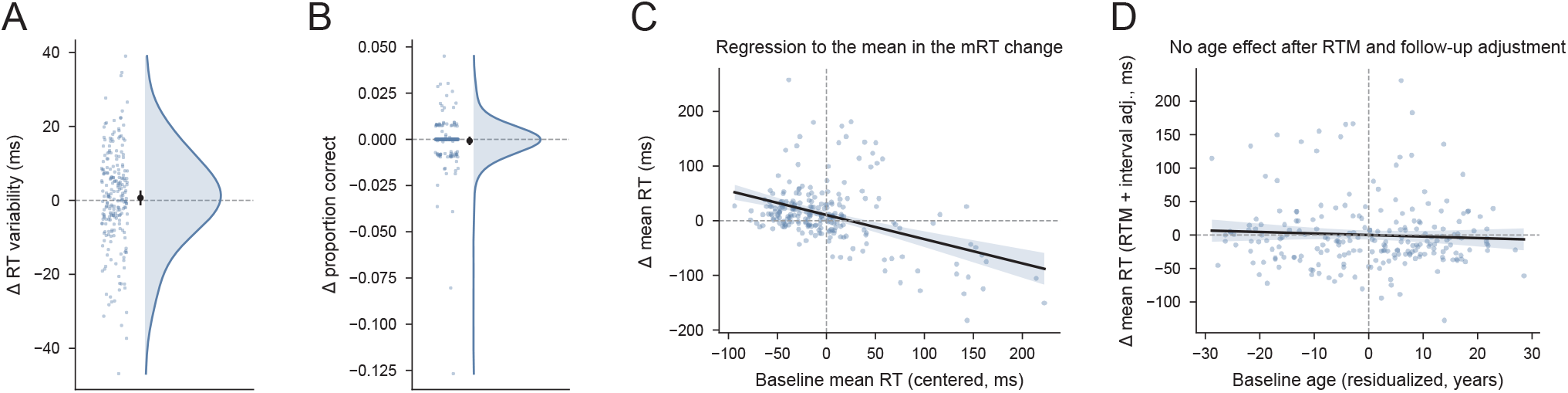
PVT reaction time variability and accuracy are unchanged, and the slowing is not an artifact of baseline dependence. (A) Within-person change in PVT reaction time variability (follow-up minus baseline). Clouds show the participant distribution and the point and whisker show the mean and its 95% confidence interval. Variability did not change (two-sided sign-flip permutation test, 10,000 permutations, p = 0.45). (B) Within-person change in response accuracy, shown as in A; accuracy did not change (two-sided sign-flip permutation test, 10,000 permutations, p = 0.38). (C) Mean reaction time change against baseline mean reaction time (centered). Slower participants at baseline tended to become faster (β = -0.44 per ms baseline, p < 0.001, adjusted R² = 0.17), but the mean increase survived this adjustment (intercept at mean baseline = 10.5 ms, p < 0.001). (D) Mean reaction time change against baseline age, with both the change and baseline age residualized on baseline performance and follow-up interval (partial-regression plot). Neither baseline age (β = -0.23 ms per year, p = 0.40) nor follow-up interval (β = 3.73 ms per year, p = 0.42) was associated with the magnitude of change. Points in C and D are individual participants; lines and shading show the ordinary least-squares fit and 95% confidence interval. n = 222 (training set) in A to D.

**Supplementary Figure 2.**
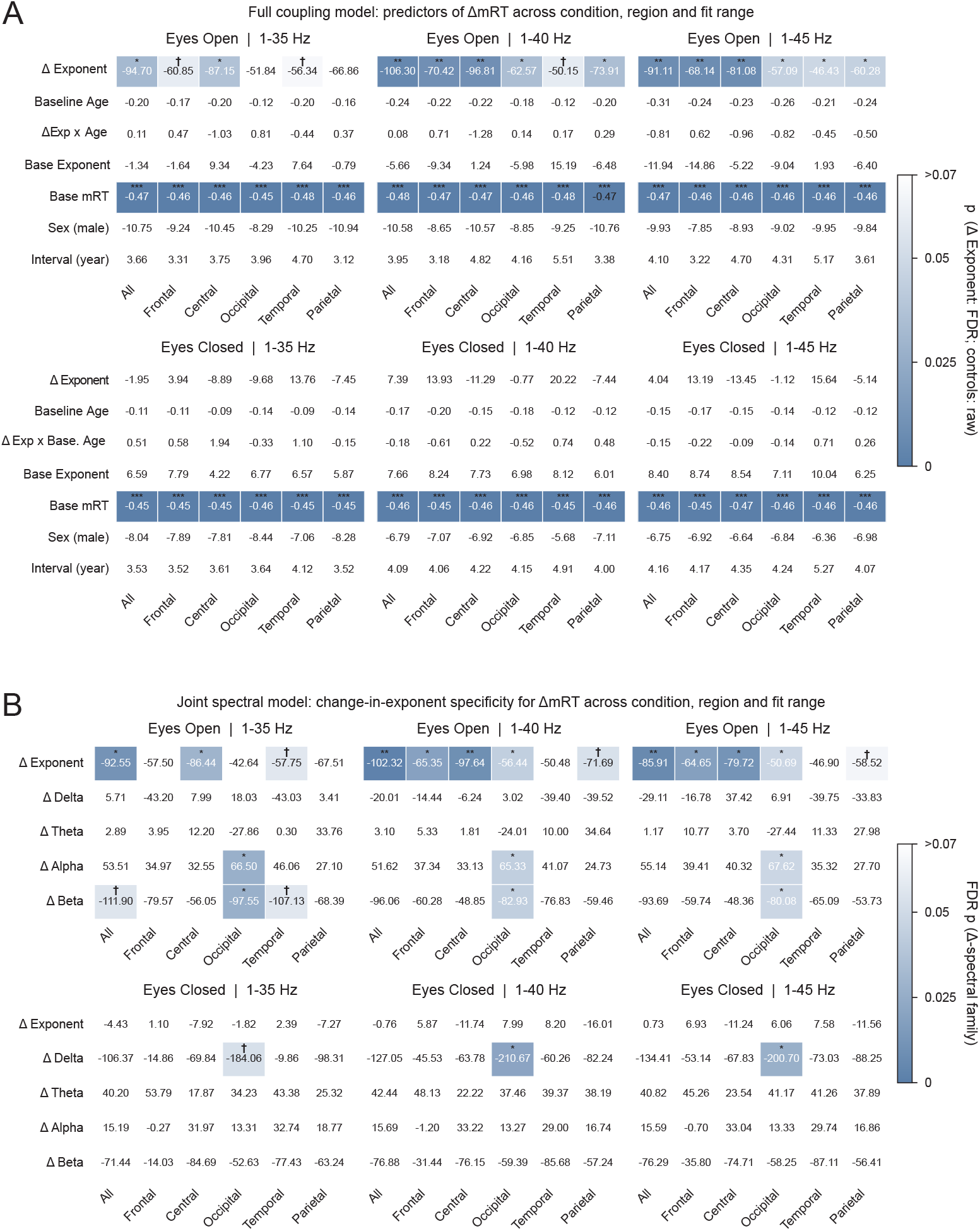
The exponent change and PVT effect is robust and largely specific across electrode regions, recording conditions and spectral parameterization fitting ranges. (A) The full coupling model of mean reaction time change (as in Fig. 3D) refitted across the six electrode regions (see Methods), the eyes-open and eyes-closed conditions, and three spectral parameterization fitting ranges (1-35, 1-40, 1-45 Hz). Each cell shows the raw coefficient and shading denotes the p-value (Exponent change β was FDR-corrected, control terms remained uncorrected). In the eyes-open condition, exponent coupling was negative in every cell and reached or approached significance across most regions and was absent throughout the eyes-closed condition. (B) The joint spectral model (exponent change with concurrent delta, theta, alpha and beta power change, as in Fig. 3E) across the same grid. Shading encodes the FDR-corrected p across the five-term spectral family. Exponent change was the most consistent and widely distributed association with slowing under eyes-open but not eyes-closed conditions. Some aperiodic-adjusted peak-power changes coupled only locally over occipital electrodes (alpha and beta under eyes-open, delta under eyes-closed). n = 222 (training set) in A and B.

**Supplementary Figure 3.**
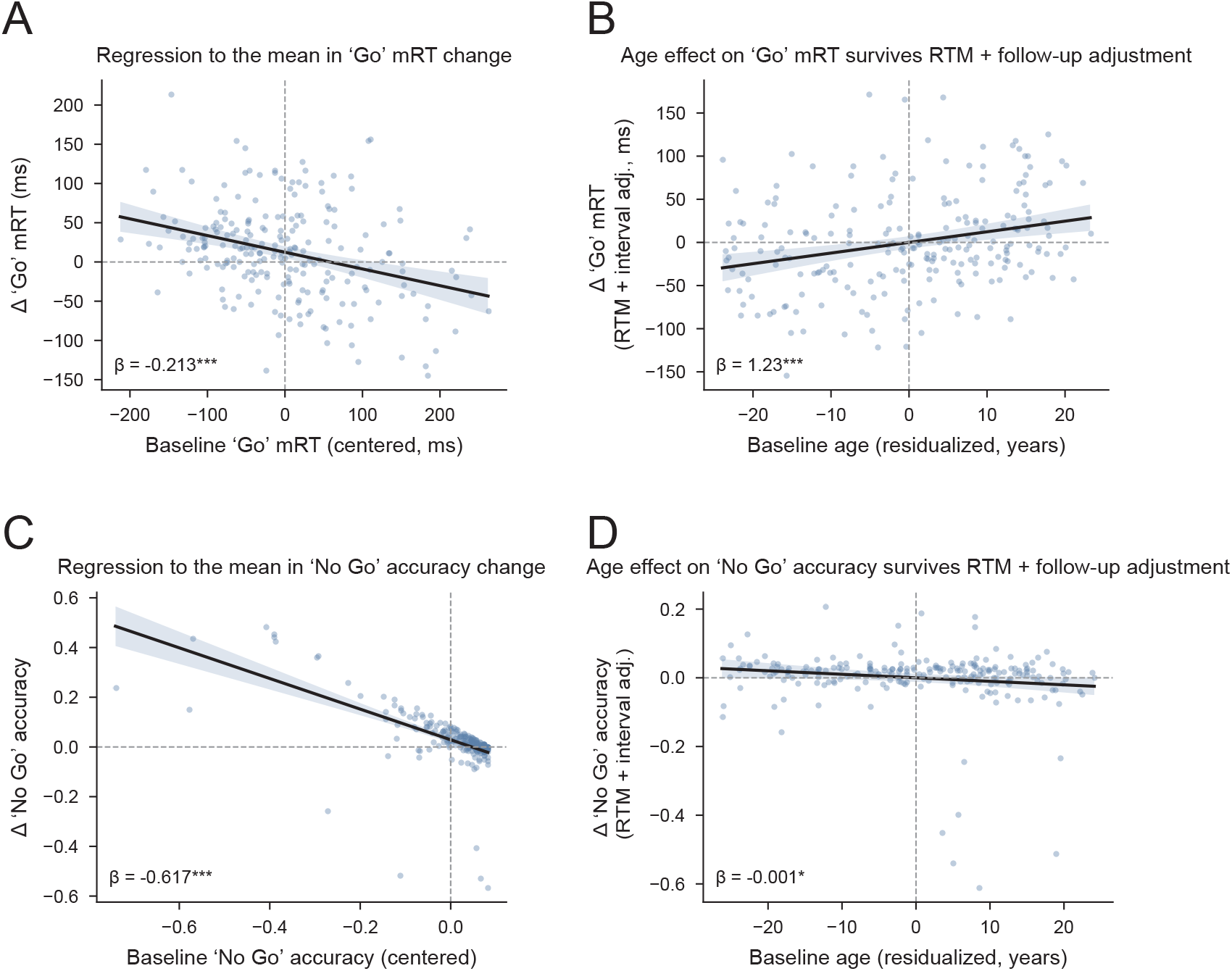
Modified Simon task response time slowing and no-go accuracy improvement survive baseline adjustment, with response-time slowing concentrated in older participants. (A) ‘Go’ response time change against baseline ‘Go’ response time (centered). Baseline level was associated with the change (β = -0.213, p < 0.001), consistent with regression to the mean, but the slowing survived this adjustment (intercept at mean baseline = 10.8 ms, p = 0.03). (B) ‘Go’ response-time change against baseline age, with both the change and baseline age residualized on baseline level, sex and follow-up interval (β = 1.23 ms per year, p < 0.001). The slowing was concentrated in older participants. (C) ‘No Go’ accuracy change against baseline ‘No Go’ accuracy (centered), shown as in A; accuracy change also showed regression to the mean (β = -0.617, p < 0.001, adjusted R² = 0.37) and survived adjustment (intercept = 0.030, p < 0.001). (D) ‘No Go’ accuracy change, residualized as in B, against baseline age. Older participants improved slightly less (age, β = -0.001 per year, p = 0.034). Points are individual participants; lines and shading show the ordinary least-squares fit and 95% confidence interval. n = 222 (training set) in A to D.

**Supplementary Figure 4.**
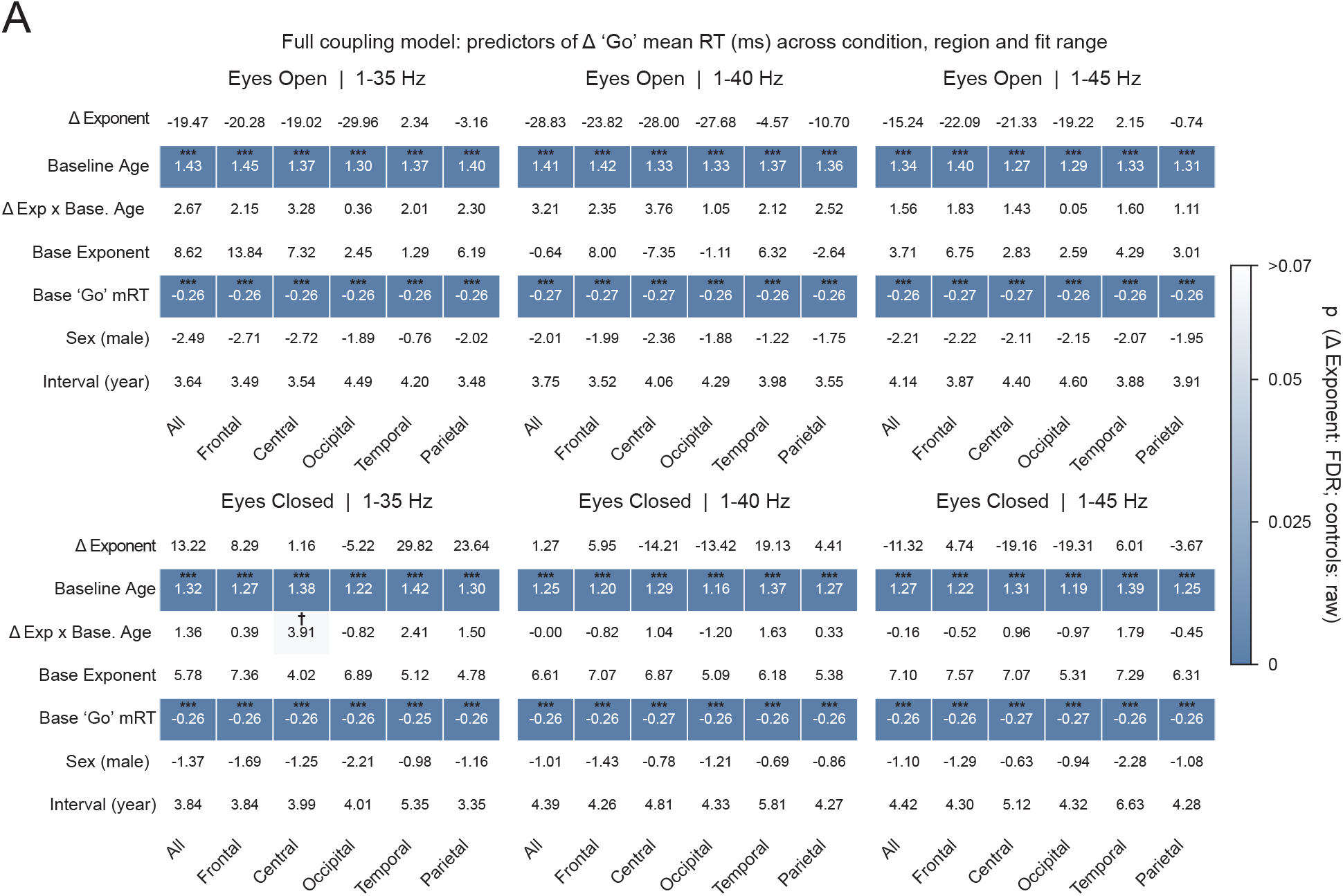
The exponent-Simon response time effect is robust across recording conditions, electrode regions and spectral parameterization fitting ranges. (A) The full coupling model of ‘Go’ mean response time change (as in Fig. 5D) refitted across the six electrode regions, the eyes-open and eyes-closed conditions, and three spectral parameterization fitting ranges (1-35, 1-40, 1-45 Hz). Each cell shows the raw coefficient and shading denotes the p-value (Exponent change β was FDR-corrected, control terms remained uncorrected). The exponent coupling did not survive correction in any region, fitting range and condition. n = 212-222 (training set) in A, depending on whether anyone was lacking a post-task eyes-open recording at one session within region.

**Supplementary Figure 5.**
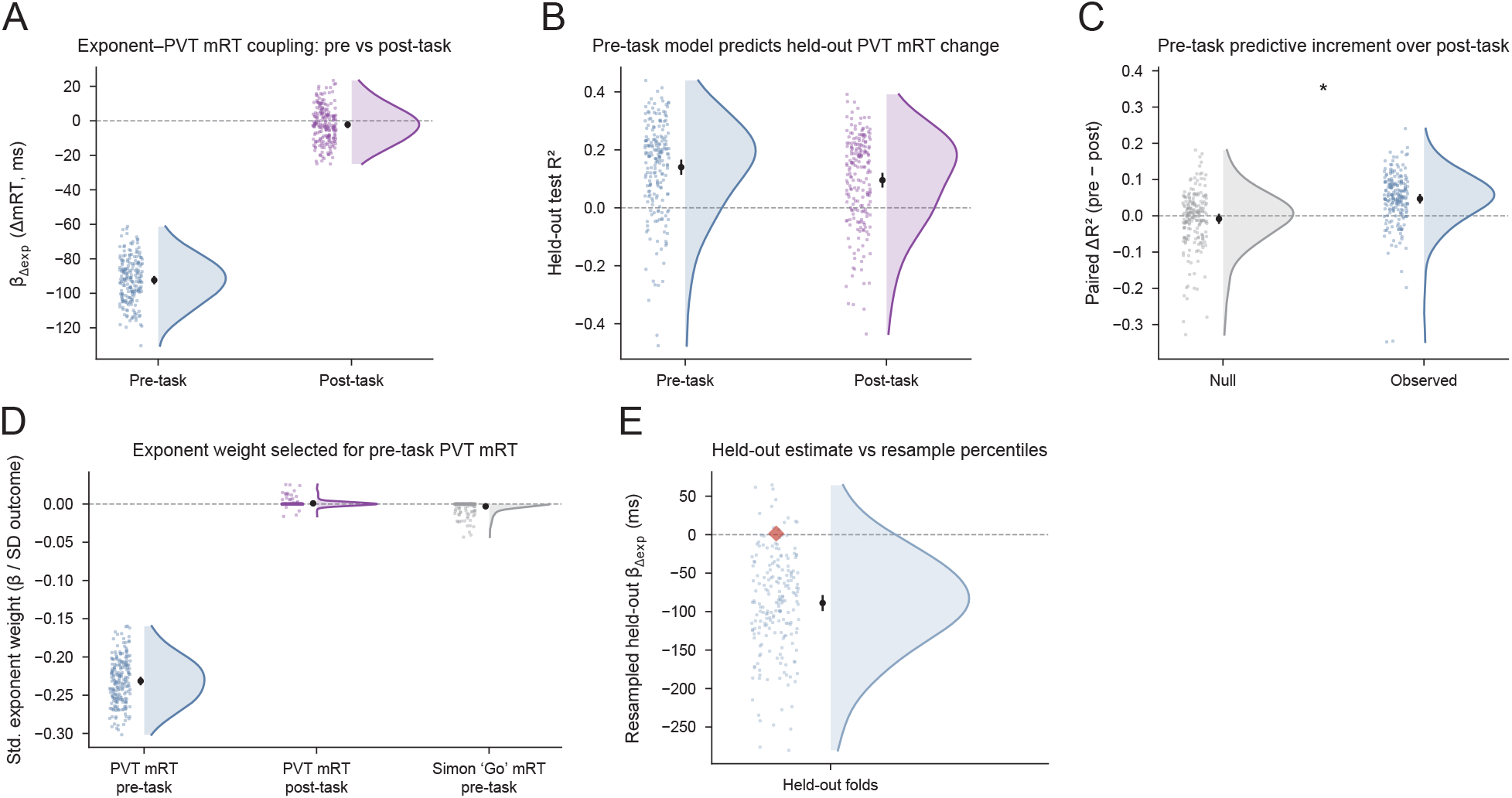
Resample stability and state- and domain-specificity of the exponent-PVT slowing coupling hold under joint age and baseline mean reaction time stratification. Conventions as in Fig. 6, with the 200 resamples stratified jointly on baseline age and baseline mean reaction time rather than baseline age alone, pre-task in blue and post-task in purple. (A) The pre-task coefficient relating exponent change to reaction-time change was negative in every resample (median = -91.88 ms) with 94% sign concordance between the independently fitted training and held-out partitions, whereas the post-task coefficient centered near zero with an unstable sign (median = -2.60 ms, negative in 59% of resamples, 21% concordance). (B) Held-out R² of each model on the held-out partition of each resample (pre-task median R² = 0.177, RMSE = 50.3 ms, positive in 81% of resamples; post-task median R² = 0.135, RMSE = 51.0 ms, positive in 73%). (C) Paired difference in held-out R² (pre-task minus post-task) on the same participants (blue) beside a null obtained by permuting the outcome within each resample’s training partition (grey). The mean paired increment exceeded a full label-shuffle null in a separate two-sided permutation test (observed mean ΔR² = 0.056; null mean = -0.002, SD = 0.019; 1,000 global permutations x 100 folds, p = 0.014); asterisks denote that permutation test. Across the 200 resamples the increment had a median of 0.052 (mean 0.047) and a 2.5th-97.5th percentile spread of [-0.090, 0.168]. (D) Standardized elastic-net exponent weight (coefficient divided by the within-fold outcome standard deviation) for PVT reaction time pre-task (blue), PVT reaction time post-task (purple) and Simon ‘Go’ response time pre-task (grey). The pre-task PVT weight was consistently negative and survived the L1 penalty in 100% of resamples (median = -0.232). The post-task PVT weight and the pre-task Simon weight were negligible (median approximately 0, retained in 12.5% and 22.5% of resamples, respectively). (E) The coupling coefficient refitted within the frozen 54-person held-out set (red diamond) against the distribution of held-out-partition coefficients from the training-set resamples (blue). The frozen coefficient was positive and non-significant (β = 1.416, p = 0.983), falling at the 94th percentile of the resample distribution, and the out-of-sample R² of the frozen training model applied to the held-out set was negative (-0.181, worse than predicting the held-out mean), at the 4.5th percentile. n = 222 (training, resampled) for pre-task panels; n = 220 for post-task panels and for all paired pre-minus-post comparisons, two participants each lacking a post-task eyes-open recording at one session; n = 54 (frozen held-out set). *p < 0.05.

**Supplementary Table 1.** Linear mixed-effects model of the aperiodic exponent and the relationship of its change to concurrent band-power change. Two models fitted to the training set (n = 222) for the whole-scalp average exponent, eyes-open condition. The upper model is a linear mixed-effects model of the exponent level, *exponent ∼ session + baseline age + sex + (1 | participant)*, fitted across both sessions (444 observations from 222 participants): Change is the within-person change across the follow-up (the binary session effect, follow-up minus baseline), baseline age (mean-centered, per year) is the between-person cross-sectional gradient, Sex (male) contrasts males with females, and the random-intercept and residual variances are the by-participant and within-participant variance components. The lower model regresses the within-person exponent change on the concurrent within-person band-power changes, *Δexponent ∼ Δdelta + Δtheta + Δalpha + Δbeta + baseline age + sex*, fitted to one change score per participant (n = 222), with each band-power change standardized (reported per standard deviation). Bands are delta (1 to 4 Hz), theta (4 to 7 Hz), alpha (7 to 13 Hz) and beta (13 to 24 Hz). β, regression coefficient in the model’s respective units; SE, standard error; CI, confidence interval; FDR p, Benjamini-Hochberg false-discovery-rate-corrected p across the four-band Δ-power family.

| Model | Term | $\beta$ | SE | 95% CI | p | FDR p | n (obs) | n (participants) |
| --- | --- | --- | --- | --- | --- | --- | --- | --- |
| **Predictors of Exponent** | Intercept | 1.097 | 0.013 | [1.071, 1.122] | < 0.001 |  | 444 | 222 |
| **(all electrodes, eyes-open)** | Change | -0.04 | 0.01 | [-0.059, -0.020] | < 0.001 |  |  |  |
|  | Baseline age (centered, /yr) | -0.005 | 0.001 | [-0.007, -0.004] | < 0.001 |  |  |  |
|  | Sex (male) | 0.026 | 0.025 | [-0.023, 0.076] | 0.295 |  |  |  |
|  | Random intercept (variance) | 0.027 |  |  |  |  |  |  |
|  | Residual (variance) | 0.011 |  |  |  |  |  |  |
| **Predictors of Exponent Change** | $\Delta$ Delta (per SD) | -0.028 | 0.011 | [-0.050, -0.006] | 0.014 | 0.056 | | |
| **(all electrodes, eyes-open)** | $\Delta$ Theta (per SD) | 0.003 | 0.011 | [-0.020, 0.025] | 0.814 | 0.814 | | |
| | $\Delta$ Alpha (per SD) | -0.004 | 0.015 | [-0.033, 0.025] | 0.776 | 0.814 | | |
| | $\Delta$ Beta (per SD) | 0.007 | 0.014 | [-0.020, 0.035] | 0.608 | 0.814 | | |
|  | Baseline age (centered, /yr) | 0 | 0.001 | [-0.002, 0.002] | 0.982 |  |  |  |
|  | Sex (male) | -0.019 | 0.021 | [-0.060, 0.022] | 0.36 |  |  |  |

**Supplementary Table 2.** Linear mixed-effects models of within-person change and between-person age effects for each cognitive-behavioral measure. Linear mixed-effects models fitted to the training set (n = 222; 444 observations from 222 participants), *measure ∼ session + baseline age + sex + (1 | participant)*, with separate models for PVT mean reaction time, Simon ‘Go’ mean reaction time, the Simon effect and Simon ‘No-Go’ accuracy. Change is the within-person change across the follow-up (the binary session effect, follow-up minus baseline); baseline age (mean-centered, per year) is the between-person cross-sectional gradient; Sex (male) contrasts males with females; the random-intercept and residual variances are the variance components. The Simon effect is defined as mean incongruent ‘Go’ reaction time minus mean congruent ‘Go’ reaction time. Reaction times are in milliseconds; ‘No-Go’ accuracy is the proportion of ‘No-Go’ trials with a correctly withheld response. β, regression coefficient; SE, standard error; CI, confidence interval. PVT, psychomotor vigilance test.

| Model | Term | $\beta$ | SE | 95% CI | p | n (obs) | n (participants) |
| --- | --- | --- | --- | --- | --- | --- | --- |
| Predictors of PVT mRT (ms) | Intercept | 266.347 | 3.774 | [258.950, 273.744] | < 0.001 | 444 | 222 |
|  | Change | 10.528 | 3.802 | [3.076, 17.979] | 0.006 |  |  |
|  | Baseline age (centered, /yr) | -0.529 | 0.249 | [-1.018, -0.040] | 0.034 |  |  |
|  | Sex (male) | -3.154 | 6.753 | [-16.389, 10.082] | 0.641 |  |  |
|  | Random intercept (variance) | 1557.682 |  |  |  |  |  |
|  | Residual (variance) | 1604.343 |  |  |  |  |  |
| Predictors of Simon Task 'Go' mRT (ms) | Intercept | 601.456 | 5.796 | [590.097, 612.816] | < 0.001 | 444 | 222 |
|  | Change | 12.397 | 4.004 | [4.549, 20.244] | 0.002 |  |  |
|  | Baseline age (centered, /yr) | 2.227 | 0.416 | [1.411, 3.042] | < 0.001 |  |  |
|  | Sex (male) | -10.778 | 11.266 | [-32.858, 11.302] | 0.339 |  |  |
|  | Random intercept (variance) | 5677.811 |  |  |  |  |  |
|  | Residual (variance) | 1779.432 |  |  |  |  |  |
| Predictors of Simon effect (ms) | Intercept | 19.26 | 1.895 | [15.546, 22.975] | < 0.001 | 444 | 222 |
|  | Change | 1.163 | 1.738 | [-2.243, 4.569] | 0.503 |  |  |
|  | Baseline age (centered, /yr) | 0.448 | 0.129 | [0.195, 0.700] | < 0.001 |  |  |
|  | Sex (male) | -4.149 | 3.489 | [-10.987, 2.689] | 0.234 |  |  |
|  | Random intercept (variance) | 462.28 |  |  |  |  |  |
|  | Residual (variance) | 335.229 |  |  |  |  |  |
| Predictors of Simon Task 'No Go' Accuracy (proportion) | Intercept | 0.918 | 0.007 | [0.904, 0.932] | < 0.001 | 444 | 222 |
|  | Change | 0.029 | 0.008 | [0.014, 0.044] | < 0.001 |  |  |
|  | Baseline age (centered, /yr) | -0.001 | 0 | [-0.002, 0.000] | 0.053 |  |  |
|  | Sex (male) | 0.001 | 0.013 | [-0.024, 0.026] | 0.932 |  |  |
|  | Random intercept (variance) | 0.005 |  |  |  |  |  |
|  | Residual (variance) | 0.007 |  |  |  |  |  |

**Supplementary Table 3.** Coupling of within-person exponent change to within-person PVT mean reaction time change, and its spectral specificity. Two ordinary least-squares regressions of the within-person change in PVT mean reaction time (follow-up minus baseline, in milliseconds) on within-person spectral change, fitted to one change score per participant from the training set (n = 222), whole-scalp average, eyes-open condition. The upper (full coupling) model is *ΔmRT ∼ Δexponent + baseline age + (Δexponent x baseline age) + baseline exponent + baseline mean RT + sex + follow-up interval*. The lower (joint spectral) model adds the concurrent change in each band’s aperiodic-adjusted peak power and their baseline levels, *ΔmRT ∼ Δexponent + Δdelta + Δtheta + Δalpha + Δbeta + (baseline level of each spectral term) + baseline age + baseline mean RT + sex + follow-up interval*. Bands denote aperiodic-adjusted peak power, the height of the largest spectral peak above the aperiodic fit within delta (1 to 4 Hz), theta (4 to 7 Hz), alpha (7 to 13 Hz) and beta (13 to 24 Hz). β, regression coefficient in milliseconds per unit predictor; SE, standard error; CI, confidence interval; raw p, uncorrected p; FDR p, Benjamini-Hochberg false-discovery-rate-corrected p (in the joint model, computed across the five-term Δ-spectral family of Δexponent plus the four bands). Baseline mean RT is mean-centered.

| Model | Term | $\beta$ | SE | 95% CI | raw p | FDR p | n (obs) | n (participants) |
| --- | --- | --- | --- | --- | --- | --- | --- | --- |
| **Predictors of change in PVT mRT (ms)**<br>(all electrodes, eyes-open) | Intercept | 148.322 | 26.267 | [96.547, 200.098] | < 0.001 |  | 444 | 222 |
| | $\Delta$ Exponent | -91.112 | 26.137 | [-142.631, -39.593] | < 0.001 | 0.008 | | |
|  | Baseline age (centered) | -0.309 | 0.298 | [-0.896, 0.278] | 0.3 |  |  |  |
| | $\Delta$ Exponent x Base. age | -0.812 | 1.629 | [-4.023, 2.400] | 0.619 | | | |
|  | Baseline exponent | -11.944 | 18.986 | [-49.368, 25.479] | 0.53 |  |  |  |
|  | Baseline mean RT | -0.467 | 0.064 | [-0.592, -0.341] | < 0.001 |  |  |  |
|  | Sex (male) | -9.93 | 7.064 | [-23.853, 3.994] | 0.161 |  |  |  |
|  | Follow-up interval (yr) | 4.096 | 4.512 | [-4.798, 12.990] | 0.365 |  |  |  |
| **Spectral Predictors of change in PVT mRT (ms)**<br>(all electrodes, eyes-open) | $\Delta$ Exponent | -85.913 | 26.105 | [-137.378, -34.447] | 0.001 | 0.006 | 444 | 222 |
| | $\Delta$ Delta SNR | -29.106 | 92.883 | [-212.225, 154.012] | 0.754 | 0.943 | | |
| | $\Delta$ Theta SNR | 1.167 | 49.901 | [-97.213, 99.547] | 0.981 | 0.981 | | |
| | $\Delta$ Alpha SNR | 55.136 | 30.552 | [-5.097, 115.368] | 0.073 | 0.121 | | |
| | $\Delta$ Beta SNR | -93.689 | 46.033 | [-184.442, -2.935] | 0.043 | 0.108 | | |
|  | Baseline exponent | -5.975 | 19.62 | [-44.654, 32.705] | 0.761 |  |  |  |
|  | Baseline Delta SNR | 95.041 | 77.61 | [-57.965, 248.048] | 0.222 |  |  |  |
|  | Baseline Theta SNR | -3.263 | 33.836 | [-69.970, 63.444] | 0.923 |  |  |  |
|  | Baseline Alpha SNR | 33.502 | 17.674 | [-1.343, 68.346] | 0.059 |  |  |  |
|  | Baseline Beta SNR | -12.767 | 22.957 | [-58.026, 32.493] | 0.579 |  |  |  |
|  | Baseline age (centered) | -0.19 | 0.297 | [-0.776, 0.395] | 0.523 |  |  |  |
|  | Baseline mean RT | -0.484 | 0.064 | [-0.611, -0.356] | < 0.001 |  |  |  |
|  | Sex (male) | -10.285 | 7.484 | [-25.040, 4.471] | 0.171 |  |  |  |
|  | Follow-up interval (yr) | 6.028 | 4.582 | [-3.005, 15.061] | 0.19 |  |  |  |

**Supplementary Table 4.** Coupling models of within-person exponent change to within-person change in each Simon-task measure. Ordinary least-squares regressions of the within-person change (follow-up minus baseline) in each Simon-task measure on within-person exponent change, fitted to one change score per participant from the training set (n = 222), whole-scalp average, eyes-open condition. Each model shares the structure of the PVT coupling model in Supplementary Table 3, *Δoutcome ∼ Δexponent + baseline age + (Δexponent x baseline age) + baseline exponent + baseline outcome + sex + follow-up interval*, with separate models for ‘Go’ mean reaction time (milliseconds), the Simon effect (milliseconds) and ‘No-Go’ accuracy (proportion). β, regression coefficient; SE, standard error; CI, confidence interval; raw p, uncorrected p; FDR p, Benjamini-Hochberg-corrected p for the Δexponent term; adj R^2^, adjusted R^2^ of the model.

| Model | Term | $\beta$ | SE | 95% CI | raw p | FDR p | n | adj R<sup>2</sup> |
| --- | --- | --- | --- | --- | --- | --- | --- | --- |
| **Predictors of Change in Simon Task 'Go' mRT (ms)**<br>(all electrodes, eyes-open) | Intercept | 167.59 | 34.574 | [99.441, 235.738] | < 0.001 |  | 222 | 0.151 |
| | $\Delta$ Exponent | -15.241 | 28.64 | [-71.693, 41.212] | 0.595 | 0.86 | | |
|  | Baseline age (centered) | 1.339 | 0.332 | [0.684, 1.994] | < 0.001 |  |  |  |
| | $\Delta$ Exponent x Base. age | 1.56 | 1.78 | [-1.948, 5.068] | 0.382 | | | |
|  | Baseline exponent | 3.706 | 20.739 | [-37.173, 44.586] | 0.858 |  |  |  |
|  | Baseline mean RT | -0.264 | 0.042 | [-0.348, -0.180] | < 0.001 |  |  |  |
|  | Sex (male) | -2.214 | 7.733 | [-17.456, 13.028] | 0.775 |  |  |  |
|  | Follow-up interval (yr) | 4.137 | 4.929 | [-5.579, 13.853] | 0.402 |  |  |  |
| **Predictors of Change in Simon effect (ms)**<br>(all electrodes, eyes-open) | Intercept | 24.375 | 8.907 | [6.818, 41.933] | 0.007 |  | 222 | 0.301 |
| | $\Delta$ Exponent | -3.102 | 11.26 | [-25.298, 19.094] | 0.783 | 0.881 | | |
|  | Baseline age (centered) | 0.068 | 0.131 | [-0.190, 0.325] | 0.605 |  |  |  |
| | $\Delta$ Exponent x Base. age | 0.705 | 0.7 | [-0.676, 2.086] | 0.315 | | | |
|  | Baseline exponent | -11.857 | 8.174 | [-27.968, 4.254] | 0.148 |  |  |  |
|  | Baseline mean RT | -0.479 | 0.049 | [-0.576, -0.381] | < 0.001 |  |  |  |
|  | Sex (male) | -2.561 | 3.043 | [-8.560, 3.437] | 0.401 |  |  |  |
|  | Follow-up interval (yr) | 1.077 | 1.957 | [-2.780, 4.935] | 0.583 |  |  |  |
| **Predictors of Change in Simon Task 'No Go' Accuracy (proportion)**<br>(all electrodes, eyes-open) | Intercept | 0.557 | 0.059 | [0.441, 0.673] | < 0.001 |  | 222 | 0.384 |
| | $\Delta$ Exponent | 0.084 | 0.047 | [-0.009, 0.177] | 0.078 | 0.7 | | |
|  | Baseline age (centered) | -0.001 | 0.001 | [-0.002, 0.000] | 0.206 |  |  |  |
| | $\Delta$ Exponent x Base. age | 0.001 | 0.003 | [-0.004, 0.007] | 0.624 | | | |
|  | Baseline exponent | 0.046 | 0.035 | [-0.022, 0.114] | 0.188 |  |  |  |
|  | Baseline mean RT | -0.627 | 0.054 | [-0.733, -0.521] | < 0.001 |  |  |  |
|  | Sex (male) | 0.003 | 0.013 | [-0.022, 0.028] | 0.827 |  |  |  |
|  | Follow-up interval (yr) | 0.009 | 0.008 | [-0.007, 0.025] | 0.276 |  |  |  |

**Supplementary Table 5.**
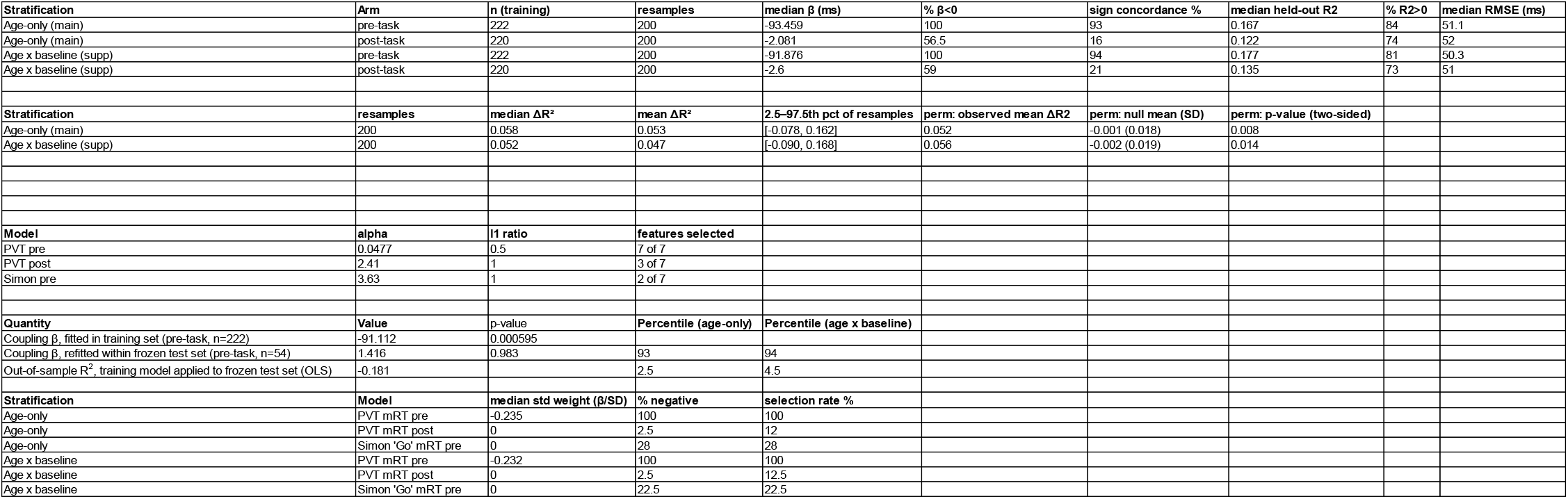
Out-of-sample predictive modelling of the exponent-PVT effect: resample stability, state specificity and domain specificity. Predictive analyses of the within-person exponent-PVT coupling across 200 age-stratified 80/20 resamples of the training set, eyes-open condition, fitted separately to pre-task and post-task EEG, with a single application of the frozen model to the held-out test set (n = 54). Pre-task models used all 222 training participants; post-task models and the paired pre-minus-post comparisons used the 220 participants with a post-task eyes-open recording at both sessions. *Out-of-sample resampling, pre- vs post-task*: n (training), the number of training participants entering each arm; the number of resamples; the median across resamples of the ordinary least-squares Δexponent coefficient (milliseconds per unit) from the coupling model; the percentage of resamples with a negative coefficient (% β < 0); sign concordance (the percentage of resamples in which the coefficient sign agreed between the independently fitted training and held-out partitions); and, scored on the held-out partition of each resample, the median R^2^, the percentage of resamples with a positive R^2^ (% R^2^ > 0), and the median RMSE in milliseconds. *Pre-task predictive increment over post-task*: the paired difference in held-out R^2^ between the pre-task and post-task models fitted on the same participants, ΔR^2^ = R^2^ (pre-task) − R^2^ (post-task), summarized across the 200 resamples as the median, the mean, and the 2.5th–97.5th percentile spread. Separately, the two-sided full label-shuffle permutation test (1,000 global permutations of the outcome x 100 folds), reported as its observed mean ΔR^2^, the mean and standard deviation of its null, and its p-value. *Tuned ElasticNet hyperparameters*: the cross-validation-tuned penalty for each ElasticNet model, alpha (overall penalty strength) and L1 ratio (the L1-to-L2 mixing parameter, searched over 0.1 to 1.0 by five-fold cross-validation), and the number of predictors retained with a non-zero weight, out of the seven coupling predictors (the same predictor set as in Table Supplement 3). *Frozen held-out split in resample context*: the coupling coefficient fitted in the training set (pre-task, n = 222) with its p-value; the coupling coefficient obtained by refitting the same model within the frozen held-out set (pre-task, n = 54) with its p-value; and the out-of-sample R^2^ obtained by applying the frozen training model to the held-out set, which is the plain coefficient of determination and is not adjusted, so a negative value indicates prediction worse than the held-out sample mean. For the two held-out quantities, the percentile column gives the position of that value within the corresponding training-resample distribution (the coefficient against the distribution of held-out-partition coefficients, the R^2^ against the distribution of held-out-partition R^2^), under each stratification. No percentile is given for the training coefficient, which is not an out-of-sample quantity. *Standardised ElasticNet weight of the exponent change*: the standardised ElasticNet weight of Δexponent, defined as the ElasticNet coefficient divided by the within-fold standard deviation of the outcome, reported as median across resamples, the percentage of resamples with a negative weight (% negative), and the selection rate (the percentage of resamples in which the L1 penalty retained a non-zero weight), for the PVT pre-task, PVT post-task and Simon ‘Go’ pre-task models under each stratification. OLS, ordinary least squares; RMSE, root-mean-squared error; SD, standard deviation; CV, cross-validation; PVT, psychomotor vigilance test.

